# CRISPR activation screen uncovers MARCKSL1 as a gauge for extracellular vesicle secretion

**DOI:** 10.1101/2025.07.24.665424

**Authors:** Yingjie Zhong, Anna E. George, Nalan Liv, Cilia de Heus, Rayman T.N. Tjokrodirijo, Manon Messchendorp, Jimmy J.L. Akkermans, Birol Cabukusta, George M.C. Janssen, Peter A. van Veelen, Ruud H.M. Wijdeven, Jacques Neefjes, Frederik Verweij, Ilana Berlin

## Abstract

Extracellular vesicles (EVs) are crucial mediators of intercellular communication that originate through one of two pathways: outward budding of the plasma membrane (PM) or fusion of mature (late) endosomes with the PM. How cells balance these EV biogenesis routes remains unclear. To address this, we performed a genome-wide CRISPR activation screen for genes that increase cell surface levels of the tetraspanin CD63—an EV marker that shuttles between late endosomes and the PM. This unbiased approach identified a membrane adaptor protein, MARCKSL1, commonly upregulated in diverse tumor types. Follow-up studies, integrating genomic activation/ablation with microscopic and proteomic approaches, revealed that MARCKSL1 potentiates EV secretion from the PM, (in part) at the expense of late endosome—PM fusion. Probing the molecular context of MARCKSL1 function, we implicate PM-bridging cytoskeletal components (e.g., Radixin) and SNARE-associated proteins (e.g., STXBP3) as collaborators of MARCKSL1. Collectively, our findings reveal new mechanistic underpinnings of PM remodeling and position MARCKSL1 as a gauge between different platforms of EV biogenesis.

## Introduction

For multicellular organisms to function coherently, individual cells contained therein must possess the means to communicate with one another. Such intercellular communication can be achieved directly via cell-cell junctions^1^ and synapses^2^ or indirectly through transmission of factors across extracellular space. The latter mode of information transfer is mediated in part by extracellular vesicles (EVs)^3^, whose secretion and uptake require complex membrane remodeling events^4^. EVs are produced by a wide variety of cell types and feature a diverse cargo repertoire of lipids, proteins and nucleic acids^5^. Collectively, these messengers potentiate (or inhibit) a wide range of fundamental biological processes, including metabolism^6^, signaling^7^, migration^8^ and epithelial-to-mesenchymal transition^9^, among many others. Correspondingly, EV-mediated communication constitutes a key aspect of physiological organ function^10,11^ and contributes to cancer progression and metastasis^12,13^ as well as dissemination of protein aggregates associated with neurodegenerative diseases^14^. Moreover, cell type-specific fusion capabilities of EVs and EV-like nanoparticles^15^ position them as promising tools for therapeutic delivery^16^ and organ repair^17,18^. Given these considerations, investigating the cellular and molecular drivers of EV biogenesis and secretion is vital to our understanding of human physiology and targeted exploitation of intercellular communication in the clinic.

EVs are subdivided into two broad classes: microvesicles (MVs) and exosomes, which are distinguished on the basis of their compartment of origin. While MVs bud outwards from the plasma membrane (PM), exosomes derive primarily from inward budding of intralumenal vesicles (ILVs) during endosomal maturation^19^. Both EV classes consist of a cytoplasmic core, bound by a lipid bilayer. Soluble factors carried by EVs include signal transducing molecules, enzymes, chaperones and various types of RNAs^20^, while their envelopes feature membrane-organizing proteins and diverse cell surface receptors^21,22^. EV-associated tetraspanins (e.g., CD63, CD81, and CD9) shuttle between the PM and endosomes and are thus enriched on EVs secreted from both compartments^23^. EV budding from the PM is stimulated by clustering of tetraspanins and other cargoes into lipid microdomains that pinch off into extracellular space, a process facilitated by localized remodeling of the actin cytoskeleton^24^. Akin to viral budding^25^, EV scission from the PM is orchestrated by the endosomal sorting complex required for transport (ESCRT)^26^. Secretion of EVs via the endocytic pathway follows similar basic principles involving cargo selection and vesicle formation but require additional fusion steps to ensure release from donor cells. Exosome biogenesis begins with sorting of endocytosed cargoes between degradation and recycling^27^. Cargoes not recycled early in the pathway become sequestered onto ILVs through ESCRT-dependent recognition^28^ or clustering activities of tetraspanins^29^, ceramides^30^, and lipid rafts^31^. The resulting late endosomes, also referred to as multivesicular bodies (MVBs), are dynamic organelles with a wide range of homeostatic and sensory functions^32,33^. At steady state, the bulk of the cell’s mature endosomes congregates in the perinuclear region^34^, where proteolytic lysosomes reside^35^. Many late endosomes go on to fuse with lysosomes^36^, committing their intralumenal cargoes for degradation, while others fuse with the PM to expel their ILVs as exosomes into extracellular space^37^. Although ILV formation is a necessary step in the biogenesis of endosome-derived EVs, it is not a terminal event, and retrofusion back to the limiting membrane provides a salvage pathway against exosome secretion^38^. To reach the PM, mature endosomes require directed transport along microtubule tracks, predicated on the small GTPase handover from Rab7 to Arl8b^39^, and ultimately Rab27a^40^, followed by productive docking and fusion. Membrane fusion is accomplished by soluble NSF-attachment protein receptors (SNAREs)^41^, in collaboration with local rearrangements of the filamentous actin (F-actin) cytoskeleton^42^. While the biogenesis of exosomes as well as microvesicles integrates membrane remodeling with actin filament dynamics, how these aspects are regulated to balance distinct EV output routes remains mysterious.

In this study, we exploited genome-wide CRISPR activation methodology^43^ to screen for new molecular players in EV biogenesis and secretion. Using cell surface levels of CD63 as a corollary for PM—endosome membrane homeostasis, we identified the membrane adaptor protein myristoylated alanine-rich C kinase substrate like protein 1 (MARCKSL1)^44^ as a positive regulator of CD63 cell surface display. Overexpression of MARCKSL1 is well documented across a wide range of cancer types^45^, and prior work has implicated its function in actin cytoskeleton remodeling and cell migration^46,47^. Additionally, MARCKSL1 has been detected in circulating EVs of colorectal cancer patients^48^, yet potential involvement of MARCKSL1 in EV biogenesis has not been investigated. By combining genetic manipulation with state-of-the art proteomic and microscopic approaches, we now reveal that upregulation of MARCKSL1 potentiates EV secretion from donor cells. Upregulation of MARCKSL1 preferentially boosts EV production from the PM (partly at the expense of MVB—PM fusion), shifting the EV cargo landscape in favor of factors involved in cell adhesion and migration. Mechanistic studies further align MARCKSL1 function with actin-associated machinery and SNARE-mediated vesicle fusion apparatus, uncovering molecular players that gauge the choice between distinct EV output routes.

## Results

### CRISPR activation screen identifies MARCKSL1 as an enhancer of CD63 cell surface display

To explore new molecular determinants of PM equilibrium and EV release, we performed a CRISPR activation screen for factors that alter cell surface display of the tetraspanin CD63. The choice of read-out was motivated by the prominence of CD63 as a marker of EVs derived from both late endosomes^49^ and the PM^50^. Human melanoma MelJuSo cells, featuring an expansive late endosomal repertoire^51^, were transduced with a CRISPR activation single-guide RNA (sgRNA) library targeting 23,000 genes^52^ and sorted for high surface expression of CD63 (Figure 1A). Subsequent gRNA enrichment analysis revealed gene products whose upregulation increased expression of CD63 at the PM (Figure 1B; Table S1). The top hit from the screen was CD63 itself, validating our chosen methodology. Additionally, we identified ACOT7 (acyl-CoA thioesterase), BCAM (basal cell adhesion molecule), COMMD7 (NFkappaB transcriptional regulator), KCNQ3 (voltage gated potassium channel), MARCKSL1, and ZNF385B (apoptotic signaling regulator) as potential modulators of CD63 membrane homeostasis (Figure 1B). Of the hits tested in the validation round, MARCKSL1 exhibited the highest effect on CD63 surface levels upon gRNA-mediated activation of its genomic locus in MelJuSo as well as HeLa cells (Figures 1C-1E, S1A and S1B). Ectopic overexpression of MARCKSL1-GFP similarly resulted in elevated surface expression of CD63 (Figures S1C-S1E).

**Figure 1.**
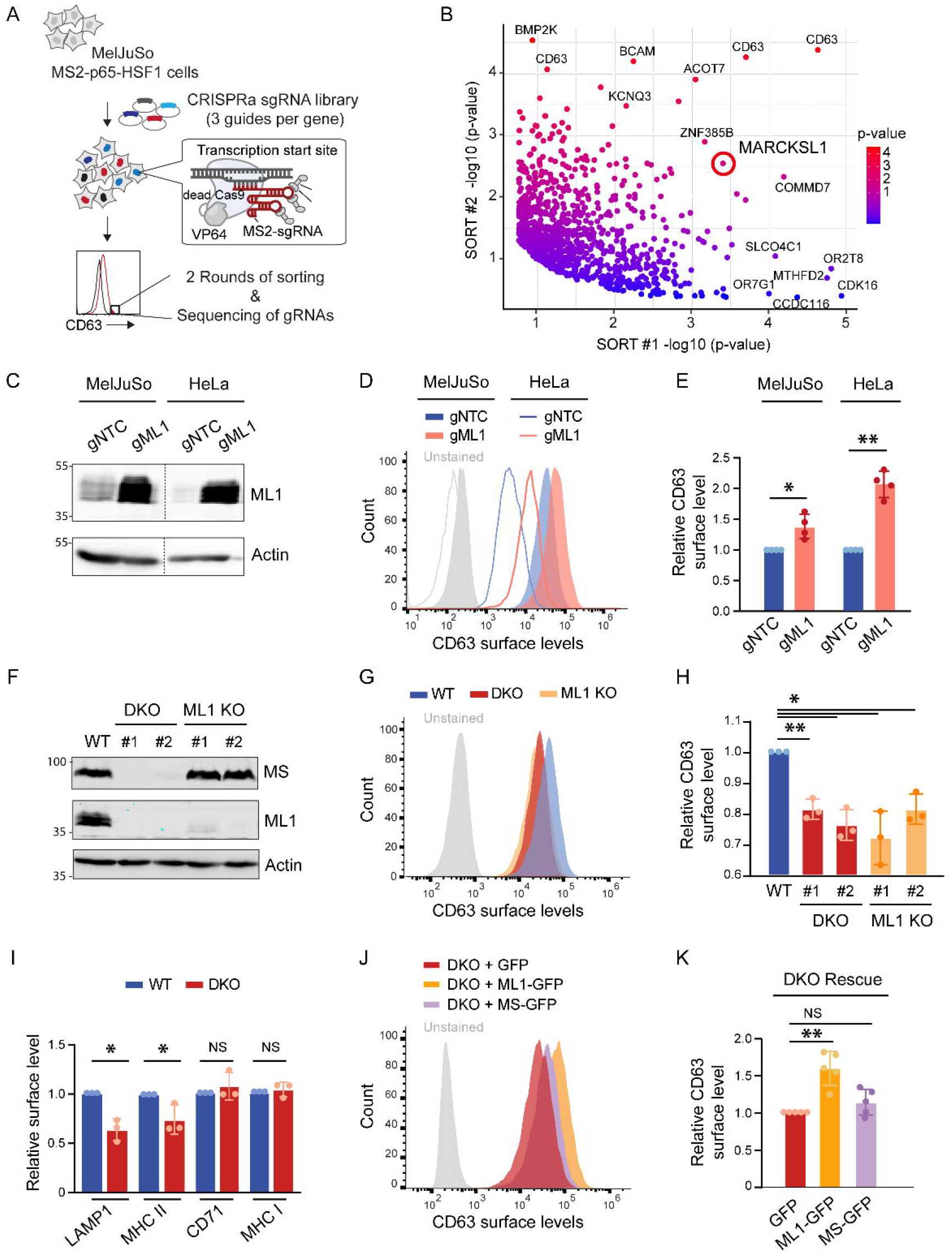
CRISPR activation screen identifies MARCKSL1 as a positive regulator of CD63 surface expression. **A**. Schematic of CRISPR activation (CRISPRa) screening methodology used in this study. MelJuSo cells stably expressing MS2-p65-HSF1 were transduced with a pooled gRNA library containing dCAS9. Cells with high CD63 surface expression were sorted twice using flow cytometry. Enriched gRNAs were identified using NGS. **B**. Scatter plot of gRNA gene targets enriched in sort 1 vs sort 2 (based on p values from RSA analysis). MARCKSL1 (highlighted) is among the top hits. See also Supplementary Figure S1A and Table S1. **C-E**. Consequences of CRISPRa-mediated MARCKSL1 upregulation for cell surface expression of CD63. **C**. Immunoblot validation of MARCKSL1 (ML1) upregulation (β-actin as loading control) in cells edited using nontargeting gRNA (gNTC) versus gRNA targeting the ML1 locus (gML1). **D**. Flow cytometry analysis of CD63 surface levels. Representative histograms of MelJuSo (filled) and HeLa (lined) cells edited as indicated are shown. **E**. Quantification of mean fluorescence intensities expressed relative to paired controls. Graph reports on n=4 independent experiments. See also Supplementary Figures S1B-S1E. **F-H**. Consequences of MARCKS/MARCKSL1 ablation for cell surface expression of CD63. **F**. Immunoblot analysis demonstrating ablation of MARCKS (MS) and/or MARCKSL1 (ML1) (β-actin as loading control). **G**. Flow cytometry analysis of wild type (WT), ML1 knock out (ML1 KO) and MS/ML1 double knock-out (DKO) MelJuSo cells. Representative histograms are shown. **H**. Quantification of mean fluorescence intensities in 2 independent clones per condition, expressed relative to WT cells. Graph reports on n=3 independent experiments. See also Supplementary Figure S2. **I**. Consequences of MARCKS/MARCKSL1 ablation for cell surface expression of endolysosomal markers LAMP1 and MHC II and biosynthetic pathway markers CD71 and MHC I. Quantification of mean fluorescence intensities based on flow cytometry analyses of DKO cells expressed relative to WT MelJuSo cells. Graph reports on n=3 independent experiments. See also Supplementary Figure S3A. **J, K**. Rescue of MARCKS/MARCKSL1 ablation. J. Flow cytometry analysis of CD63 surface levels in MelJuSo DKO cells stably transduced with empty vector (GFP), MS-GFP or ML1-GFP. Representative histograms are shown. **K**. Quantification of mean fluorescence intensities expressed relative to GFP-expressing DKO cells. Graph reports on n=5 independent experiments. See also Supplementary Figures S3B-S3E. All bar graphs reflect mean (+/-SD) with statistical significance determined using paired Student *t*-test. *p < 0.05, **p < 0.01, NS: not significant.

To investigate the role of MARCKSL1 in CD63^+^ membrane homeostasis, we performed loss-of-function and rescue studies in MARCKSL1 knock-out cells. Because MelJuSo and HeLa cells also express MARCKS – a close relative of MARCKSL1 – we went on to create double knock-out cells (DKO) lacking both gene products (Figures 1F and S2). Flow cytometry analysis demonstrated significant reductions in CD63 levels at the surface of ML1 KO and DKO cells, as compared to their parental (wild type, WT) counterparts (Figures 1G and 1H). Cell surface expression of other markers shuttling between late endosomes and the PM, such as LAMP1 and MHC class II (major histocompatibility complex class II receptor), was also reduced in DKO cells (Figures 1I and S3A). By contrast, presence of housekeeping receptors CD71 and MHC class I at the surface of DKO cells remained unaffected (Figures 1I and S3A). Comparable phenotype severity exhibited by ML1 KO and DKO cells (Figures 1F-H) suggested that MARCKS may not serve an important role here. This was confirmed by stable re-expression of MARCKS1, which efficiently rescued surface expression of CD63 (and LAMP1) in DKO cells, while MARCKS afforded to significant benefit under the same conditions (Figures 1J, 1K, and S3B-S3E). We therefore concluded that loss of MARCKSL1 cannot be compensated by MARCKS in this context.

### MARCKSL1 potentiates EV secretion

Alterations in protein composition at the PM, particularly with respect to tetraspanins like CD63, have direct implications for EV biogenesis^53^. Having identified MARCKSL1 as a positive regulator of CD63 residence at the PM, we proceeded to examine the contribution of MARCKSL1 to EV secretion. To this end, EV isolates from culture media of parental versus ML1 KO or DKO MelJuSo cells were analyzed using nanoparticle tracking analysis (NTA) and immunoblotting^38,54^ (Figure 2A). NTA measurements indicated a reduction in the number of particles isolated from DKO cells as compared to the control, without appreciably affecting particle size distribution (Figures 2B and S3F). In line with this, immunoblot analysis of EV fractions revealed reduced abundance of CD63, CD81 and MHC II relative to the corresponding whole cell lysate (WCL) inputs (Figures 2C, 2D, S3G and S3H). Importantly, no appreciable effects on total cellular levels of EV protein markers were observed (Figure 2E). In contrast to ablation, upregulation of endogenous MARCKSL1 (gML1) boosted EV release relative to control (gNTC), as evidenced by both nanoparticle profiling and immunoblotting (Figures 3A and 3B). Here, markedly elevated levels of CD63 and MHC II were detected in EV fractions isolated from cells overexpressing endogenous MARCKSL1, while total protein expression of these markers remained unaffected (Figures 3B and 3C). Increase in EV-associated CD81 appeared less prominent (Figures 3B and 3C), suggesting a potential cargo shift in EVs induced by cells that upregulate MARCKSL1.

**Figure 2.**
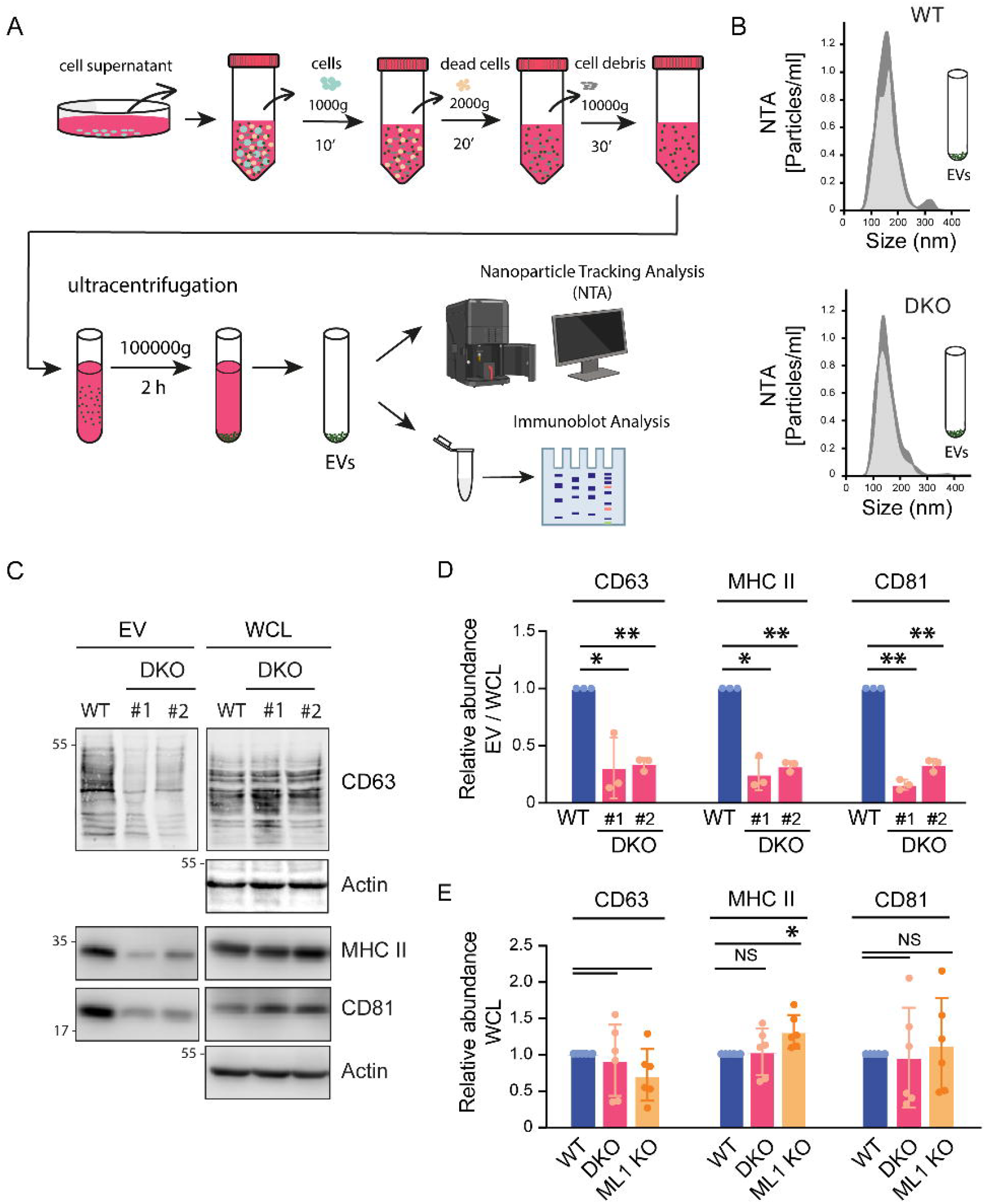
MARCKS/MARCKSL1 ablation reduces EV secretion. **A**. Schematic representation of EV isolation and analysis workflow used in this study. **B-E**. Consequences of MARCKS/MARCKSL1 ablation for EV release. **B**. Particle size profiling of EV isolates from wild type (WT) versus double knock-out (DKO) MelJuSo cells using Nanoparticle Tracking Analysis (NTA). Representative histogram plots of particle size distribution (nm) are shown, with thick gray lines reflecting deviation between injection rounds. **C**. Immunoblot analysis of EV isolates and their corresponding whole cell lysate (WCL) controls derived from WT and DKO cells for EV markers CD63, CD81 and MHC II. **D**. Quantification of protein abundance in EV isolates expressed as ratio EV / WCL. Graph reports on n=3 independent experiments. **E**. Quantification of total protein abundance for EV and endolysosomal markers CD81, LAMP1 and MHC II in DKO and ML1 KO cells expressed relative to WT MelJuSo cells. Graph reports on n=6 independent experiments. See also Supplementary Figures S3F-S3H. All bar graphs reflect mean (+/-SD), with statistical significance determined using paired Student *t*-test. *p < 0.05, **p < 0.01, NS: not significant.

**Figure 3.**
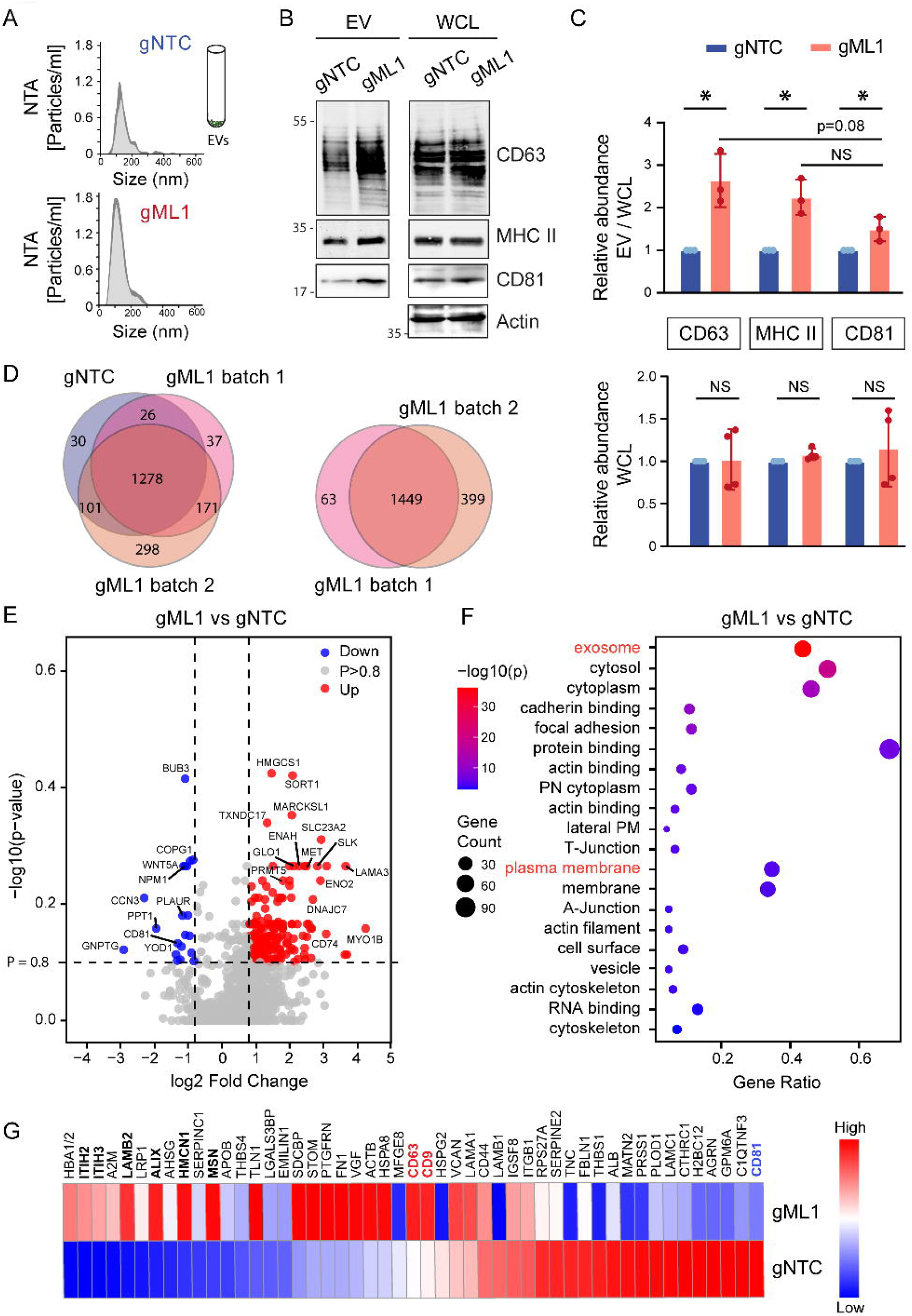
Upregulation of MARCKSL1 boosts EV secretion. **A-C**. Impact of MARCKSL1 upregulation on EV biogenesis. **A**. Particle size profiling of EV isolates from control (gNTC) versus endogenous MARCKSL1 overexpressing (gML1) MelJuSo cells using NTA. Representative histogram plots of particle size distribution (nm) are shown, with thick gray lines reflecting deviation between injection rounds. **B**. Immunoblot analysis of EV isolates and their corresponding whole cell lysate (WCL) controls for EV markers CD63 and MHC II. **C**. Quantification of protein abundance in EVs expressed as ratio EV / WCL (top) and WCL only (bottom) as a function of MARCKSL1 upregulation. Bar graphs reflect mean (+/-SD) of n=4 independent experiments, with statistical significance determined using paired Student *t*-test; *p < 0.05, NS: not significant. **D-G**. Proteomic profiling of EVs released from MelJuSo cells in response to MARCKSL1 upregulation. **D**. VEN diagrams comparing the average of n=2 independent gNTC batches (blue) with gML1 batches 1 (pink) and 2 (coral) and gML1 batches to each other. **E**. Volcano plot highlighting proteins enriched in (p>0.8) (red) or depleted from (blue) EV isolates of gML1 vs. gNTC cells. **F**. Gene ontology (GO) clustering of proteins enriched in EVs isolated from gML1 vs. gNTC cells. **G**. Heat map comparing relative abundance (gML1/gNTC) for the top 50 proteins overall identified in EVs isolates from both groups. Tetraspanins whose relative abundance is up (red) or down (blue) in both gML1 replicates vs. gNTC are marked. Other differentially abundant proteins of interest are highlighted (bold font). See also Supplementary Table S2.

To profile the cargo repertoire on EVs triggered by MARCKSL1, we performed mass spectrometry analysis of EV isolates derived from the media of gNTC versus gML1 MelJuSo cells. Roughly 1500 proteins were identified per condition (Figure 3D, Table S2) using previously described EV purification methodology^55^. Although the EV proteome of cells activated for MARCKSL1 exhibited broad overlap with that of control cells, certain cargoes were enriched (Figures 3E-3G). Gene ontology (GO) clustering revealed strong association with the PM and extracellular exosome categories (p value < 0,01) (Figure 3F). In line with prior characterization of EVs from colorectal cancer patients^48^, MARCKSL1 itself was readily detected in EV isolates and was among the top differentially abundant proteins identified between gNTC and gML1 samples (Figures 3E). Other proteins involved in cell adhesion and migration (e.g., ENAH, LAMA3, and MYO1B) were also enriched in EVs derived from gML1 cells (Figure 3E). Of the 50 most abundant EV cargoes, extracellular matrix stabilizers ITIH2/3, membrane remodeler PDCD6IP (also known as ALIX), and cytoskeleton-associated proteins LAMB2 and MSN were detected in higher quantities in gML1 samples relative to control (Figure 3G). CD63 was also modestly enriched in gML1 samples (Figure 3G), which may have been due to its improved cell surface availability under MARCKSL1 overexpression (Figure 1). Another EV-associated tetraspanin, CD9, was similarly enriched in gML1 samples, while CD81 was depleted in comparison (Figure 3G). Extracellular matrix proteins Matrilin-2 (MATN2), Thrombospondin-1 (THBS1) and Tenascin-C (TNC) were also less abundant in EVs derived from gML1 cells (Figure 3G), while presence of certain enzymes (e.g., ENO2, GLO1, HMGCS1, PRMT5, SLK, and TXNDC17) was enhanced in the same samples (Figure 3E). Taken together with the loss and gain of function studies described above, these experiments support a positive role for MARCKSL1 in EV biogenesis, with implications for cargo incorporation.

### MARCKSL1 regulates trafficking of CD63 to the PM

Given that exosomes and microvesicles harbor partially overlapping cargoes, we proceeded to investigate which EV biogenesis pathways are triggered by MARCKSL1. Using CRISPR/Cas-mediated genome editing to knock-in a GFP fluorophore into the endogenous MARCKSL1 locus, we observed accumulation of MARCKSL1 at the PM and on late endosomal membranes (Figures 4A and S4A; Movie S1), suggesting that it could stimulate EV production from both compartments. MARCKSL1 contains three conserved structural elements: an N-terminal myristylation site, a MARCKS Homology 2 (MH2) domain, and an effector domain (ED) featuring phosphorylation sitesz. In line with previous findings^44^, effector domain removal (Δ ED-GFP) rendered MARCKSL1 unable to localize at the PM but had no negative effect on its localization to late endosomes (Figures 4A and S4B). Although dynamic membrane association of MARCKSL1 was reported previously^44^, no functional relevance for this protein in the context of endocytosis or extracellular vesicle biology has been described. We found that MARCKSL1 lacking its effector domain (but not the MH2 domain) was unable to upregulate CD63 on the surface of affected cells and failed to rescue reduced surface expression of CD63 in DKO cells (Figures 4B, S4C and S4D). These observations indicated that association with the PM enables MARCKSL1 to regulate the distribution of CD63. To test the impact of MARCKSL1 on CD63 trafficking between the PM and endosomes, we followed its endocytosis and recycling using pulse-chase antibody labeling experiments. Constitutive internalization of CD63 from the PM was found to be enhanced in DKO cells relative to wild type cells (Figures 4C and 4D), while recycling of CD63 to the cell surface was reduced under the same conditions (Figures 4E and 4F). Collectively, these changes likely account for the loss of CD63 surface expression incurred by MARCKS/MARCKSL1 ablation.

**Figure 4.**
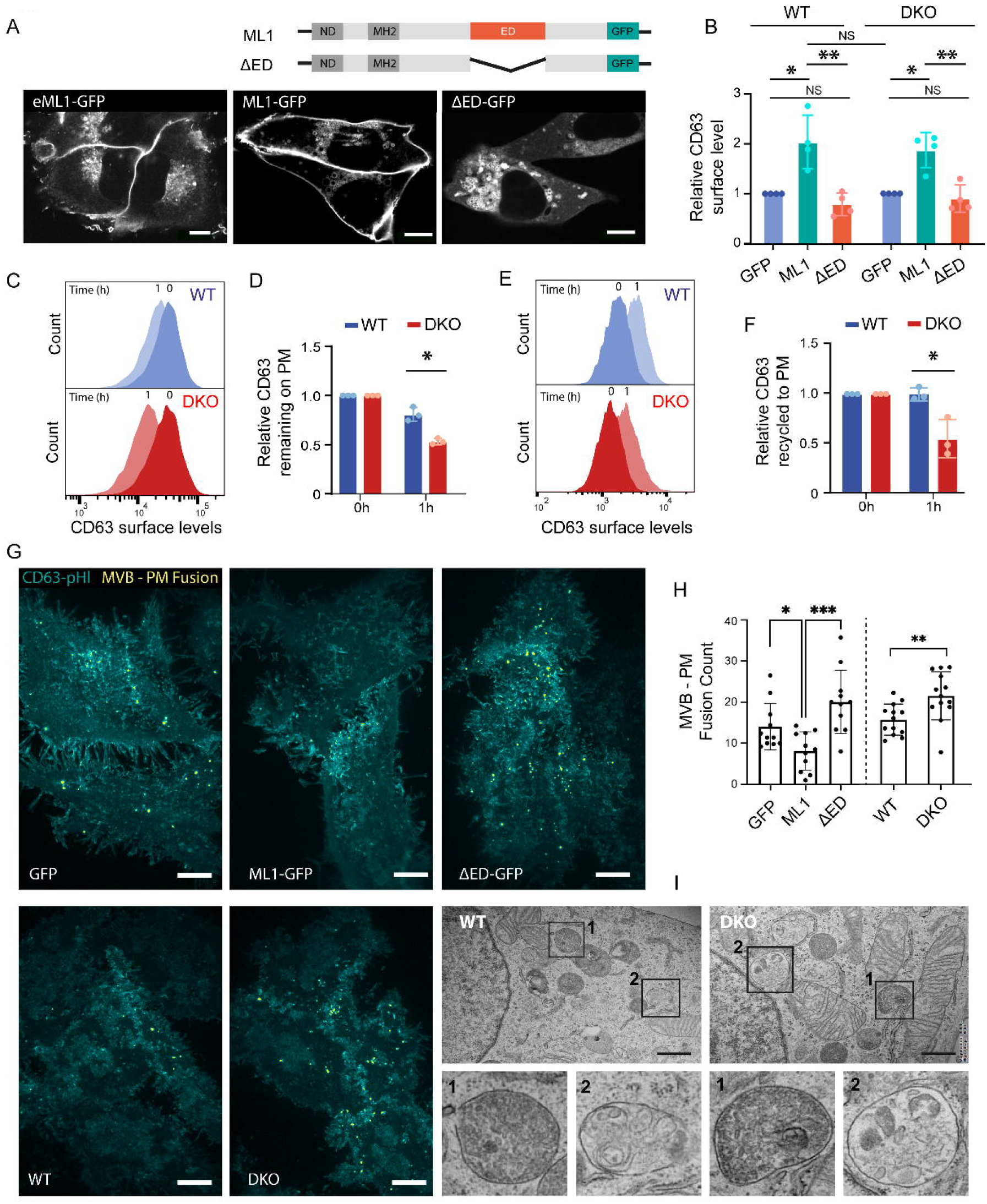
Membrane adaptor MARCKSL1 regulates CD63 trafficking but does not promote MVB— PM fusion. **A**. Top: Schematic representation of wild type MARCKSL1 domain organization (ND: N-terminal myristylation site, MH2: MARCKS Homology 2 domain, ED: effector domain) and effector domain deletion (Δ ED) mutant. Bottom: Effect of ED on the intracellular distribution of MARCKSL1. Representative confocal image of live MelJuSo cells expressing either endogenous (eML1-GFP) or transfected fluorescent MARCKSL1 (ML1-GFP versus Δ ED-GFP) constructs. Scale bars = 10 um. See also Supplementary Figures S4A and S4B and Movie S1. **B**. Effect of ED on MARCKSL1-mediated regulation of CD63 cell surface expression. Quantification of mean fluorescence intensities expressed relative to control cells (GFP) within each group. Graph reports on n=4 independent experiments. See also Supplementary Figures S4C and S4D. **C-F**. Effect of MARCKS/MARCKSL1 ablation on endocytosis and recycling of CD63. **C**. Representative histogram plots and **D**. quantification of CD63 remaining at the cell surface as a function of time (h) in wild type (WT) versus double knock-out (DKO) MelJuSo cells. **E**. Representative histogram plots and **F**. quantification of CD63 recycled to the cell surface as a function of time (h) in WT versus DKO cells. Bar graphs report mean fluorescence intensities of CD63 at the cell surface, expressed relative to WT cells for each time point, from n=3 independent experiments, with statistical significance determined using unpaired Student *t*-test. **G, H**. Effects of MARCKSL1 expression on MVB—PM fusion. **G**. Representative maximum projections of CD63-pHluorin signal (cyan) and fusion events (yellow) from representative TIRF time-lapse acquisitions. Top panels: comparison of MelJuSo cells ectopically expressing GFP, ML1-GFP or Δ ED-GFP. Bottom panels: comparison of WT versus DKO MelJuSo cells. Scale bars = 10 um. **H**. Quantification of CD63-pHluorin^+^ fusion events. Graphs report on n=3 independent experiments. See also Supplementary movies S2-S6. **I**. Effects of MARCKS/MARCKSL1 ablation on the ultrastructure of endolysosomes. Representative electron micrographs (EM) of WT and DKO MelJuSo cells are shown. Zoom-ins highlight organelles with densely packed ILVs (1) and endosomes with heterogeneous luminal content (2). Scale bars = 500 nm. All bar graphs reflect mean (+/-SD), with statistical significance determined using paired Student *t*-test; *p < 0.05, **p < 0.01, **p < 0.001, NS: not significant.

To explore how the above considerations may relate to different EV biogenesis pathways, we exploited the CD63-pHluorin reporter system for monitoring individual MVB – PM fusion events in real time^56,57^. This assay takes advantage of fluorophore quenching in the acidic environment of late endosomes, which is rapidly restored upon fusion with the PM^58^. Using this approach, we found that transient overexpression of MARCKSL1-GFP did not enhance MVB – PM fusion counts, instead slightly reducing their occurrence relative to vector control (Figures 4G and 4H; Movies S2A and S2B). This reduction was in turn abolished upon ED removal from MARCKSL1 (ξED-GFP) (Figures 4G and 4H; Movie S2C). Conversely, MARCKS/MARCKSL1 ablation exhibited the opposite effect, resulting in modestly elevated fusion counts (Figures 4G and 4H; Movies S2D and S2E). However, biogenesis of mature endosomes – the principal intracellular membrane reservoirs of CD63 – remained unaffected, as MVBs in cells lacking MARCKS/MARCKSL1 exhibited normal intraluminal contents (Figure 4I). These observations indicated that upregulation of MARCKSL1 suppresses endosome-derived EV secretion, instead boosting EV release from the PM.

### STXBP3 and Radixin collaborate with MARCKSL1 to control CD63 cell surface display

To probe the molecular mechanism(s) underlying MARCKSL1 function in PM homeostasis and EV release, we explored its molecular context inside the cell using proximity-based biotinylation methodology^59^. Mass spectrometric analysis of neutravidin precipitates from cells ectopically expressing TurboID (TID) fusions of MARCKS (MS-TID) or MARCKSL1 (ML1-TID), and treated with exogenous biotin, identified 749 proteins with unique peptides compared to empty vector expressing cells (Figures 5A and S5A; Table S3). Gene ontology clustering of candidate interactors yielded 28 significantly enriched categories, notably including extracellular exosome and apical and lateral membrane GO terms (Figure 5B). STRING network analysis of 40 highest ranked candidates revealed clusters corresponding to actin filament binding, calmodulin binding, membrane binding and docking, solute transport across membranes, and vesicle transport (Figure 5C). Among the top 10 hits were the cytoskeletal adaptor protein Radixin (RDX), known to link cortical actin filaments to the PM^60^; unconventional PM-associated myosin motor Myo1D^61^; EH domain-containing protein 4 (EHD4) involved in membrane tubulation^62^, and syntaxin-binding proteins 2 and 3 (STXBP2/3) implicated in SNARE-mediated membrane fusion^63^ (Figure 5B, 5C). Time-resolved validation of several candidate partners using biotinylation, followed by immunoblotting, confirmed intracellular proximity of MARCKSL1 to Radixin, STXBP3 and EHD4 (Figures 5D, 5E and S5B-S5E). Moesin (MSN), another member of the EMR (Ezrin, Moesin, Radixin) family^64^, was also identified with proximity-based biotinylation (Figure 5C) and found to be enriched in EVs purified from MARCKSL1 overexpressing cells (Figure 3G), suggesting that interactions at the interface between the actin cytoskeleton and the PM constitute an important feature of MARCKSL1 function in the context of EV biology.

**Figure 5.**
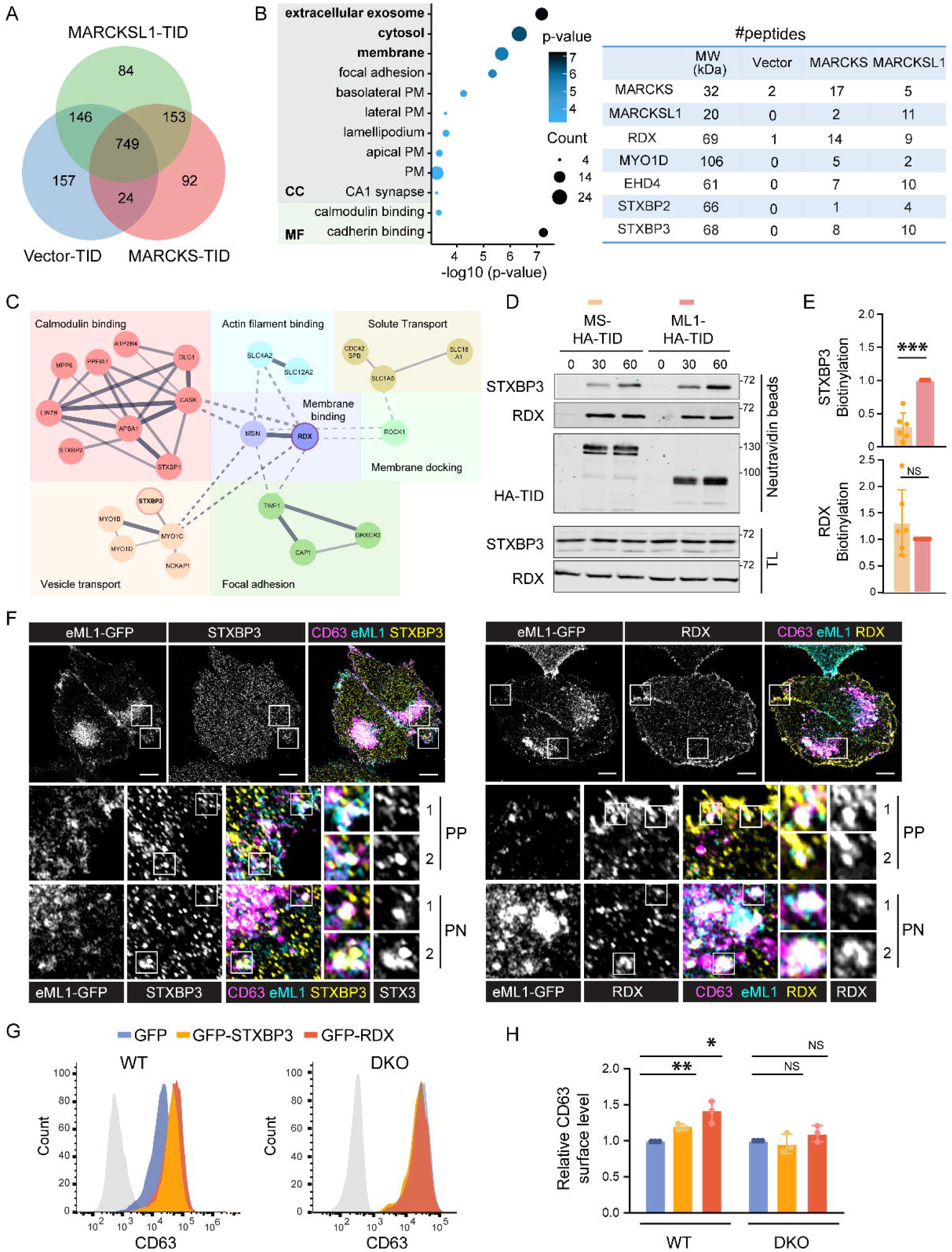
Proximity-based proteomics for MARCKSL1 identify PM anchor proteins and membrane fusion machinery. **A-C**. Proximity-based interactome of MARCKS/MARCKSL1. **A**. VEN diagram of proteins identified by proteomic analysis of neutravidin precipitates derived from HEK293T cells transfected with 2HA-TurboID empty vector (TID, blue), MARCKS-TID (pink) or MARCKSL1-TID (green) and treated in the presence of exogenous biotin. **B**. Left: GO term enrichment for 749 candidate interactors of MARCKS and MARCKSL1 (above vector control). Right: list of top 5 hits from (A). **C**. STRING analysis of 25 highest confidence hits for MARCKSL1, clustered based on common cellular functions (each group individually colored). Interactions between and within groups are indicated with dashed and solid lines, respectively. See also Supplementary Figure S5A and Table S3. **D-F**. Validation of candidate interactors of MARCKS (MS) and MARCKSL1 (ML1). **D**. Immunoblot analysis of neutravidin precipitates and their corresponding total lysate (TL) inputs from HEK293T cells transfected with MS-HA-TID or ML1-HA-TID and treated with exogenous biotin for the indicated time (min) prior to lysis. **E**. Quantification of biotinylation (relative to ML1). Graph reports on n=3 independent experiments. **F**. Colocalization of MARCKSL1 with STXBP3 and Radixin (RDX). Representative confocal images of fixed MelJuSo cells expressing endogenous ML1-GFP (cyan), immunostained for CD63 (magenta) and either STXBP3 or RDX (yellow). Zoom-ins highlight perinuclear (PN) and peripheral (PP) cell regions and areas of colocalization between channels. Scale bars = 10 um. See also Supplementary Figures S5B-S5E. **G, H**. Effects of STXBP3 and Radixin overexpression on CD63 membrane homeostasis. **G**. Representative flow cytometry histograms of CD63 surface expression in wild type (WT) versus double knock-out (DKO) MelJuSo cells, ectopically expressing GFP, GFP-STXBP3, or GFP-RDX. **H**. Quantification of CD63 surface levels, expressed relative to GFP within each group. Graph reports on n=3 independent experiments. See also Supplementary Figures S5F and S5G. All bar graphs reflect mean (+/-SD) with statistical significance determined using a paired Student *t*-test; *p < 0.05, **p < 0.01, ***p < 0.001, NS: not significant.

To explore whether MARCKSL1 collaborators contribute to PM homeostasis and EV biogenesis, we tested their intracellular localization and effects on cell surface expression of CD63. Endogenous STXBP3 distributed predominantly to intracellular puncta, some of which were found to colocalize with MARCKSL1 and CD63 (Figure 5F). Like MARCKSL1, endogenous Radixin was strongly enriched at the PM, though colocalization between the two proteins with CD63^+^ vesicles could also be observed (Figure 5F). Overexpression of either GFP-STXBP3 or GFP-Radixin phenocopied upregulation of MARCKSL1, resulting in elevated levels of CD63 at the cell surface (Figures 5G, 5H, S5F and S5G). Importantly, these effects were no longer observed in DKO cells (Figures 5G, 5H), implying that modulation of CD63 membrane homeostasis by STXBP3 and Radixin proteins relies on MARCKS/ MARCKSL1.

Taken together, the evidence presented in this study supports a model wherein MARCKSL1 enhances cell surface residence of CD63, thereby shifting the balance of EV secretion towards the PM route (Figure 6). In this capacity, MARCKSL1 collaborates with proteins at the interface between the PM and the actin cytoskeleton (e.g., Radixin and MSN), as well as those associated with the vesicle fusion apparatus (i.e., STXBP3). Our observations thus provide new molecular insights into the regulation of tetraspanin homeostasis, with implications for EV biogenesis and related membrane processes.

**Figure 6.**
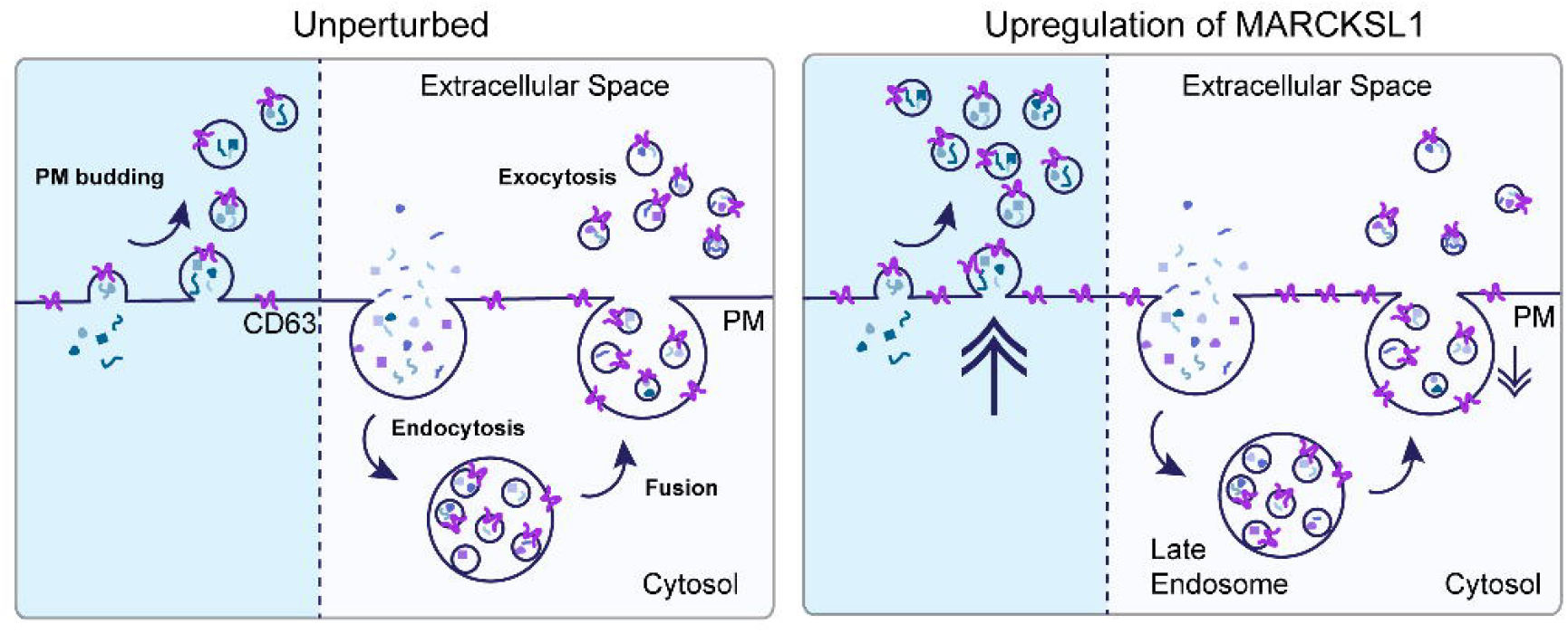
MARCKSL1 expression gauges the balance between PM-based and endosomal platforms for EV biogenesis. The evidence presented in this study supports a model wherein MARCKSL1 enhances cell surface residence of CD63, thereby shifting the balance between EV biogenesis routes. In cells upregulating MARCKSL1 expression, PM-derived EV secretion is elevated, whereas fusion of late endosomes with the PM is suppressed.

## Discussion

Controlled communication between cells is paramount to effective cross-tissue coordination and organismal homeostasis. In recent years, extracellular vesicles have emerged as potent factors in human development, health and disease. To elucidate relevant pathophysiological mechanisms and enable therapeutic cooption of EV-mediated material transfer, a detailed molecular understanding of EV release and targeting is required. While EVs carry a wide variety of cargoes, tetraspanins, including CD63, count among their most common features. Taking advantage of this, we employed CRISPR activation methodology to screen for genetic triggers of EV secretion through the prism of altered CD63 membrane homeostasis. This unbiased approach identified MARCKSL1 as a promotive factor in EV biogenesis, revealing a new player in extracellular communication with potential implications for EV-based technology development.

The MARCKS family of proteins has been implicated in embryonic development an organogenesis^65^. Additionally, MARCKSL1 is upregulated in a broad range of cancers and associated with poor prognosis^45^. Of relevance to the work presented here, MARCKSL1 was detected in circulating EVs of colorectal cancer patients^48^, albeit its potential involvement in membrane dynamics or EV secretion have not been previously explored. We found that upregulation of the genomic locus encoding MARCKSL1 leads to elevated levels of CD63 on the surface of affected cells, while its loss exerts the opposite effect. Importantly, a closely related family member, MARCKS, is unable to compensate for loss of MARCKSL1 with respect to CD63 trafficking, underscoring functional diversity. It has recently become appreciated that accumulation of tetraspanins, such as CD63, at the PM provides a driving force for microvesicle budding and shifts the balance between endosome dependent and independent EV biogenesis pathways^50^. In line with this, we found that overexpression of MARCKSL1, a membrane adaptor^66^ enriched at the PM and on endolysosomes^44^, suppresses MVB—PM fusion, while simultaneously greatly enhancing overall EV secretion. Our findings thus expand the emerging paradigm of give-and-take between distinct EV secretion platforms and position MARCKSL1 as a molecular gauge for EV output.

Prior studies have demonstrates that MARCKSL1 stabilizes cortical actin filaments^46^ in the context of cell migration^67^. Using proteomic profiling of EV isolates and proximity-based interaction studies we have identified actin bridging ERM proteins (e.g., Radixin and Moesin) as likely collaborators of MARCKSL1 at the interface between cortical actin filaments and the PM. ERM proteins function as subcellular scaffolds by anchoring transmembrane proteins to the cytoskeleton and, like MARCKSL1, promote cell migration in the realm of organ development and cancer^68^. Considering the importance of local actin filaments as a source of force during EV budding from the PM^24^, partnerships with ERM proteins may enable MARCKSL1 to effectively integrate cytoskeletal dynamics with membrane remodeling events. In addition to proteins associated with the cytoskeleton, we also identified SNARE-binding components as interactors of MARCKSL1. Among these, STXBP3, also known as Munc18c, has recently been shown to accelerate SNARE-dependent membrane fusion^69,70^. Physiologically, STXBP3 is implicated in docking and fusion of intracellular GLUT4-containing vesicles with the cell surface in adipocytes and, in so doing, regulate glucose-stimulated insulin secretion^71^. Moreover, loss of STXBP3 has been shown to induce scattering of recycling Rab11 vesicles throughout the cytoplasm, leading to diminished transport in intestinal epithelial cells^72^. These considerations suggest that, by engaging different intracellular partners, MARCKSL1 could tip the balance between vesicle budding and fusion at the PM, and in so doing, influence the EV cargo landscape.

Collectively, our findings provide new insights into the molecular underpinnings of intercellular communication through EV secretion. Given the emerging recognition of EVs as pro-tumorigenic mediators^12^, unraveling the molecular drivers of EV release carries therapeutic potential. Prior efforts of others have laid the groundwork for exploiting MARCKSL1 in cancer diagnostics and therapy. In this context, our current findings position MARCKSL1 as a candidate for targeted protein degradation^73,74^, with removal of MARCKSL1 expected to hamper cell-cell crosstalk and pathogenic transmission via EVs. At the same time, upregulation of MARCKSL1 could be envisioned as a tool to boost and EV production.

## Materials and Methods

### Antibodies and reagents

#### Flow cytometry

Primary mouse anti-CD63 NKI-C3 (1:100), CD71 (1:100, Biolegend), LAMP1 (H4A3) (1:100, Santa Cruz Biotechnology), MHC I (1:100, Biolegend), and MHC II (1:100, Biolegend) were used, followed by anti-mouse Alexa-dye-coupled secondary antibodies (1:200, Invitrogen).

#### Immunoblotting

The following primary antibodies were used: mouse anti-CD63 NKI-C3 (1:1000), mouse anti-MHC II (1:1000, NKI-1B5), mouse anti-CD81 (1:1000, Santa Cruz Biotechnology), LAMP1 (H4A3) (1:1000, Santa Cruz Biotechnology), mouse anti-actin (1:3000, Sigma-Aldrich, A5441), mouse anti-HA (1:1000, Covance, 16B12), mouse anti-FLAG (1:1000, Sigma-Aldrich, M2), rabbit anti-STXBP3 (1:1000, ThermoFisher), and rabbit anti-Radixin(1:1000, abcam). The following secondary antibodies were used: rabbit anti-mouse-HRP, sheep anti-rabbit-HRP (Invitrogen), goat anti-rabbit and goat anti-mouse IRdye 680 (1:20,000) or IRdye 800 (1:10,000) (LiCor).

#### Confocal microscopy

For immunofluorescence, mouse anti-CD63 NKI-C3 (1:500), rabbit anti-STXBP3 (1:500, ThermoFisher), rabbit anti-Radixin (1:100, abcam), and rabbit anti-EHD4 (1:100, ThermoFisher) were used, followed by Alexa-dye-coupled secondary antibodies (1:400, Invitrogen). SiR-lysosome (1:2,000, Spirochrome) or LysoTracker Red (Life Technologies, 1:10,000) were used to label mature endosomes and lysosomes in live cells.

### Cell lines, culture and transfections

MelJuSo (human melanoma) cells were cultured in IMDM (GIBCO). HeLa (human cervical carcinoma) and HEK293T (human embryonic kidney) cells were cultured in DMEM (GIBCO), all supplemented with 8% fetal calf serum (FCS, Greiner). For exosome isolation, cells were maintained in 8% exosome-depleted fetal bovine serum (ThermoFisher Scientific). All cell lines were cultured at 37°C and 5% CO2 and routinely tested (negatively) for mycoplasma. MelJuSo cells were authenticated using Sanger sequencing. HeLa and HEK293T cells were obtained from ATCC.

For transient transfection, cells were seeded in culture plates to reach ∼ 70% confluency on the day of transfection. MelJuSo cells were transfected with XtremeGene (2,5ug DNA per well in 6-well plates or 35mm dishes, 0.5 µg DNA per well of 24-well plates) according to the manufacturer’s protocol. HeLa and HEK293T cells were transfected using polyethylenimine (PEI, Polysciences) at a ratio of 3⍰μg PEI per μg DNA in 200⍰μl DMEM. Transfection complexes were allowed to form for 15⍰ min at RT and added dropwise to the cells. Treated cells were cultured overnight prior to analysis.

### DNA Constructs

MARCKS and MARCKSL1 (and mutants) were expressed from C1 CMV promoter vector series and cloned into mGFP-N1 by EcoRI/BamH1 restriction sites. Turbo-MARCKS/L1 were subcloned between EcoRI and BamHI sites of the N1-Turbo-2HA vector using standard PCR methods. Expression constructs for MARCKS and MARCKSL1 were made using Gateway cloning of the indicated genes into lentiCRISPR v2 vector (a gift from Feng Zhang, Addgene #49535), and pLenti-CAG-gate-FLAG-IRES-GFP vector (a gift from William Kaelin, Addgene #107398). All constructs were sequence verified.

### CRISPR activation screen

CRISPR activation screen was performed according to previously described methodology^52^ adapted to using CD63 cell surface labeling as a readout, as follows. Human CRISPR two-plasmid Synergistic Activation Mediator pooled library (SAM v2) used for the screen was a gift from Feng Zhang (Addgene #1000000078)^75^. 150 million MelJuSo cells stably expressing MS2-p65-HSF1 (MPH)^52^ activator were transduced at a multiplicity of infection of 0.3 with the guide RNA plasmid library. The next day, cells were selected with hygromycin (200 µg/ml) and blasticidin (2.5 µg/ml). After 5 days, two batches of cells were immunostained for CD63 and top 10% of CD63-expressing cells were sorted out using flow cytometry (sort 1). Cells were then allowed to recover and grow out for an additional 6 days. Cells were sorted once again (sort 2) to select high CD63 expressors. For each sorted cohort, cells were grown to 10 million and genomic DNA was isolated and amplified using established protocols using established methodology^75^. gRNAs were sequenced using the Illumina HighSeq 2500, and inserts were mapped to the reference. Statistical analysis was performed using redundant siRNA activity (RSA) analysis, with enrichment >4 considered a hit^76^.

### CRISPR/Cas9 genome editing of MARCKS/MARCKLS1

For activation of MARCKSL1, non-targeting gRNA sequence and gRNA targeting the genomic locus of MARCKSL1 (primer sequences in Table I) were cloned into the into BsmBI site of lentiSAMv2 vector (Addgene #75112), containing coding sequences for dCas9 and blasticidin resistance. To generate viral particles, HEK293T cells were transfected with packaging plasmids psPAX2 and pCMV-VSVG in combination with the lentiviral construct. Virus was harvested, filtered, and target cells were transduced in the presence of 8 µg/ml Polybrene (Millipore). For gene ablation, gRNA sequences targeting the genomic loci of MARCKS and/or MARCKSL1 (primer sequences in Table I) were cloned into the BbsI site of PX440 vector (containing coding sequences for Cas9 and puromycin resistance).

**Table I.**
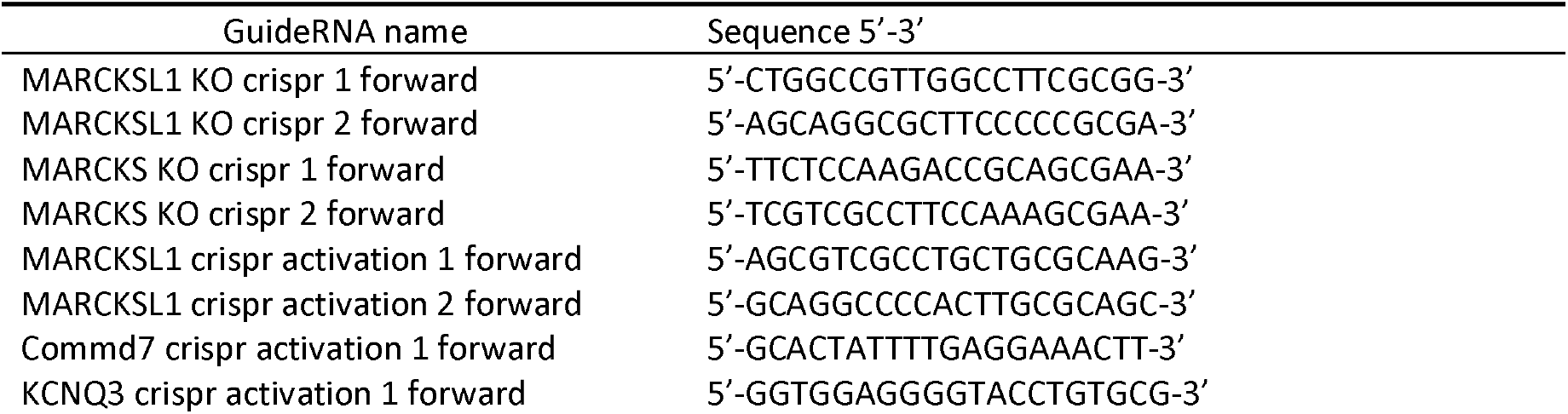
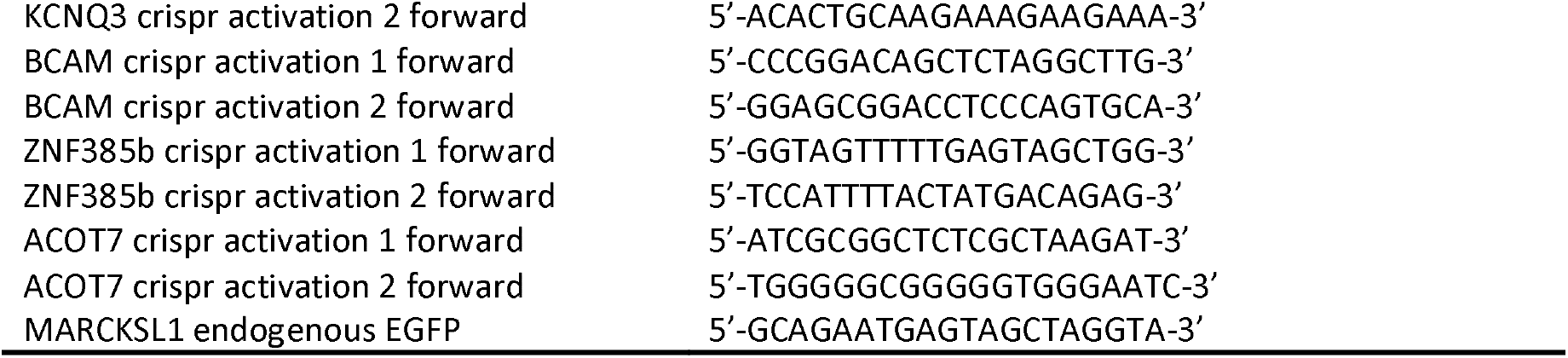
GuideRNA sequences.

Generation of MelJuSo cells expressing endogenous MARCKSL1-GFP was performed using knock-in methodology, as previously described^77^. Briefly, mGFP was derived from mGFP-C1 (Clontech) construct, wherein the promoter and pA terminator were replaced by noncoding regions from PUC19 construct. mGFP cojoined by a T2A and antibiotic selection marker was inserted, flanked by a multiple cloning site on either side. gRNA primer sequences targeting the genomic locus of MARCKSL1 are listed in Tables I and II. Constructs in question were transfected into MelJuSo cells and, the following day, cells were selected with 1 µg/ml puromycin for 3 days. Then, cells were reseeded into a 15 cm dish, allowing colonies to grow from single cells. Clonal lines were isolated, expanded and analyzed for loss/gain/tagging of MARCKS/L1 using genomic sequencing and immunoblotting.

**Table II.**
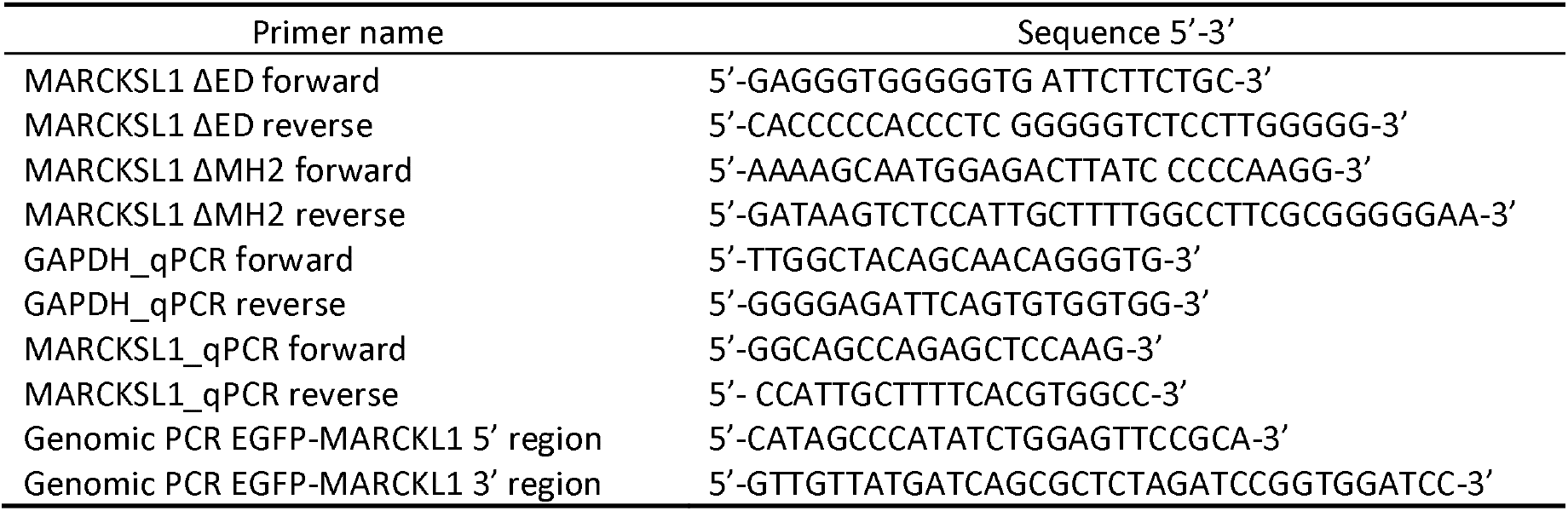
Primer sequences.

### Flow cytometry

Cells were grown in 24-well plates O/N, washed with PBS, trypsinized and stained using primary Abs against the indicated markers in 2% FCS/PBS for 30 min, followed by secondary Abs for 25 min at 4,^°^C. Cells were then washed and transferred to 1,4 mL polystyrene Round-Bottom Tubes (Micronic) and surface fluorescence levels were measured on the BD LSR II. Data were analyzed using FlowJo, with geometric mean of fluorescence intensity used to draw comparisons between samples.

### EV isolation and nanoparticle tracking analysis (NTA)

Cells were incubated in exosome-depleted medium for 24h. Supernatants (enriched in EVs) were retrieved and subjected to sequential differential centrifugation at 4°C as follows: 1000 g for 10 min; 2000 g for 20 min; 10000 g for 30 min. Cells were collected separately and counted to ensure that EV isolate comparisons were based on the same number of cells. Post-10000 g supernatants were transferred to ultra-tube and centrifuged further at 100,000 × g for 2h.

For NTA, EV isolates were diluted in PBS filtered through 0.22-µm filters to ensure appropriate particle detection (10–100) per frame. Captures and quantification were performed using the built-in NanoSight Software v3.2. Sample measurements were performed at 21°C, with the camera level set at 7 as all small particles are visible while reducing background noise. For each measurement, samples were introduced into the chamber using 1 ml syringes (Fisher Scientific), and three consecutive 60-s recordings were acquired at 25 frames per second and analyzed with a detection threshold 3 to determine EV concentration and size. Analysis of EV fractions was carried out using NanoSight NS300 (Malvern Panalytical Ltd., Malvern, UK). EV samples were further processed for immunoblot analysis by solubilization in buffer containing 0.5% NP-40, 150 mM NaCl, 50 mM Tris-HCl pH 7.6 and 5 mM MgCl2 for 30 min at 4°C prior to NTA analysis.

For mass spectrometry, EV samples were prepared as previously described^55^. Briefly, proteins were concentrated using trichloroacetic acid (TCA) precipitation. Sodium deoxycholate was added to the EV samples with a concentration of 2 mg/mL and ice-cold 100% (w/v) TCA (Sigma-Aldrich, 91230) was added to samples with a concentration of 20% and mixed immediately. Samples were incubated for 30 minutes at 4°C and centrifuged at 16,000 g for 10 minutes at 4°C. Pellets were washed twice with 1 mL of pre-cooled acetone, centrifuged as above and air dried prior to analysis (details provided under Mass Spectrometry).

### Proximity-based biotinylation interactomics

To probe the interactome of MARCKS/MARCKL1 we employed proximity-based biotinylation using the TurboID (TID) module^59^. HEK293T cells were transfected with the indicated constructs harboring 2xHA-TID modules, and biotinylation was carried out 18h post-transfection. Biotin (Sigma) was dissolved directly into serum-containing cell culture medium to a final concentration of 50 μM and added to cells. Treated cells were returned to the incubator for 30 or 60 min, as indicated, allowing for biotinylation to take place. Cells were then washed twice with PBS and lysed in TID lysis buffer (0.5% NP-40, 50 mM Tris-HCl pH 8.0, 150 mM NaCl, 5 mM MgCl, 1% SDS, protease inhibitors) to stop the labeling. Lysates were incubated rotating for 20 min at 4°C and clarified for 20 min at 12,000 rpm. Biotinylated proteins were precipitated from cell lysates using Pierce™ High Capacity NeutrAvidin™ Agarose beads (Thermo Fisher Scientific), incubated rotating at 4°C overnight. Beads were washed 4x in TID lysis buffer and samples were prepared by denaturation in Laemmli Sample Buffer. Proteins were separated by SDSPAGE and analyzed by mass spectrometry (details provided under Mass Spectrometry) or immunoblotting.

### Mass spectrometry

For MS analysis of TurboID samples, gel bands were reduced with 10 mM dithiothreitol, alkylated with 55 mM iodoacetamide. In-gel trypsin digestion was performed using a Proteineer DP digestion robot (Bruker). Tryptic peptides were extracted from the gel slices using 50/50/0.1 water/acetonitrile /formic acid (v/v/v). Samples were then lyophilized and stored at -20 °C until further processing. Samples were dissolved in water/formic acid (100/0.1 v/v) and subsequently analyzed by on-line C18 nanoHPLC MS/MS with a system consisting of an Ultimate3000nano gradient HPLC system (Thermo, Bremen, Germany), and an Exploris480 mass spectrometer (Thermo) as in^78^. Fractions were injected onto a cartridge precolumn (300 μm × 5 mm, C18 PepMap, 5 um, 100 A, and eluted via a homemade analytical nano-HPLC column (50 cm × 75 μm; Reprosil-Pur C18-AQ 1.9 um, 120 A (Dr. Maisch, Ammerbuch, Germany). The gradient was run from 2% to 40% solvent B (20/80/0.1 water/acetonitrile/formic acid (FA) v/v) in 30 min. The nano-HPLC column was drawn to a tip of ∼10 μm and acted as the electrospray needle of the MS source. The mass spectrometer was operated in data-dependent MS/MS mode for a cycle time of 3 seconds, with a HCD collision energy at 30% and recording of the MS2 spectrum in the orbitrap, with a quadrupole isolation width of 0.8 Da. In the master scan (MS1) the resolution was 120,000, the scan range 400-1500, at standard AGC target @maximum fill time of 50 ms. A lock mass correction on the background ion m/z=445.12 was used. Precursors were dynamically excluded after n=1 with an exclusion duration of 10 s, with a precursor range of 20 ppm. Charge states 2-5 were included. For MS2 the first mass was set to 110 Da, and the MS2 scan resolution was 30,000 at an AGC target of 100% @maximum fill time of 60 ms. In post-analysis processing, raw data were first converted to peak lists using Proteome Discoverer version 2.4 (Thermo Electron) and subsequently submitted to the Uniprot Homo sapiens minimal database (20205 entries), using Mascot v. 2.2.04 (www.matrixscience.com) for protein identification. Mascot searches were performed with 10 ppm and 0.02 Da deviation for precursor and fragment mass, respectively and the enzyme trypsin was specified. Up to three missed cleavages were allowed. Oxidation on Met and acetylation on protein N-term were set as a variable modification; carbamidomethyl on Cys was set as a fixed modification. Peptides with an FDR<1% were accepted.

For MS analysis of EV TCA precipitates, lysis was performed using 5% SDS lysis buffer (100 mM Tris-HCl pH7.6) and 5 U benzonase nuclease (Thermo Scientific) with incubation at 95 °C for 4 min. Protein concentrations were determined using Pierce BCA Gold protein assay (Thermo Fisher Scientific). 100 µg protein of each sample was then reduced with 5 mM TCEP, and reduced disulfide bonds were subsequently alkylated using 15 mM iodoacetamide. Excess iodoacetamide was quenched using 10 mM DTT. Protein lysates were precipitated using chloroform/methanol; resulting pellets were re-solubilized in 40 mM HEPES pH 8.4 and digested using TPCK treated trypsin (1:12.5 enzyme/protein ratio) overnight at 37⍰°C. Peptide concentration was then determined using Pierce BCA Gold protein assay. Subsequently, samples were cleaned up by solid phase extraction in 1 ml 0.1% formic acid and eluted using 1cc C18 SPE cartridges (Oasis HLB, Waters) using 400 ul of 35% acetonitrile in 0.1% formic acid. Samples were then lyophilized and stored at -20 °C until further processing. Prior to measurement, samples were dissolved in water/formic acid (100/0.1 v/v) and analyzed by on-line C18 nanoHPLC MS/MS with a system consisting of a Vanquish Neo and an Orbitrap Astral mass spectrometer (Thermo). The Vanquish Neo was operated in trap and elute mode. Samples were injected onto a cartridge precolumn (300 μm × 5 mm, C18 PepMap, 5 um, 100 A) and eluted via a homemade analytical nano-HPLC column (30 cm × 75 μm; Reprosil-Pur C18-AQ 1.9 um, 120 A (Dr. Maisch, Ammerbuch, Germany) at a flow of 350 nl/min. The gradient was run from 2% to 40% solvent B (20/80/0.1 water/acetonitrile/formic acid (FA) v/v) in 20 min. The nano-HPLC column was drawn to a tip of ∼10 μm and acted as the electrospray needle of the MS source. The mass spectrometer was operated in DIA mode at a resolution of 240,000 in MS1, a scan range of 380-980 Th at an AGC value of 500% and a maximum fill time of 3 ms. MS2 was recorded at an HCD collision energy of 25%, with a 2 Th isolation window, and a maximum fill time of 5 ms with and recording of the MS2 spectrum in the Astral. A lock mass correction on the background ion m/z=202.0777 was used.

DIA raw data were processed using Proteome Discoverer version 3.2 (Thermo Electron), using the Chimerys mode with inferys 4.7.0 as prediction model, and with default settings. In short, the settings were: trypsin as the enzyme (max 2 missed cleavages), peptide length between 7 and 30, charge between 1 and 4, and a fragment tolerance of 20 mmu. Oxidation on methionine was set as a dynamic modification and carbamidometyl on cysteine as a fixed modification. FDR was set to 1%. Quantitation was done in MS Apex mode. The minimal human database (20,596 entries) was used.

### Confocal fluorescence microscopy

For live cell imaging, cells were grown in live cell dishes (35mm) O/N, transfected, if applicable, for 24 h and incubated with the indicated cell-permeable dyes. Images and time-lapses were acquired on a Dragonfly 200 spinning disc microscope equipped with a Sona camera (Andor), solid state lasers and a climate control chamber using 63× oil immersion objectives in a 2,048 × 2,048 format.

For immunofluorescence, cells were grown on coverslips (Menzel Glaser) in 12-well plates for 24 h, fixed (PBS/3.7% formaldehyde 15 min at RT; 3 x PBS washes), permeabilized (PBS/0.1% TritonX-100 10 min at RT; 1 x PBS wash), and immunostained using the indicated primary antibodies in blocking buffer (PBS/0.5% bovine serum albumin (BSA, A8022, Sigma) 1h at RT), followed by the appropriate secondary antibodies as previously described^79^. Coverslips were mounted on glass slides (Fisher Scientific) using ProLong Gold mounting medium with DAPI (Life Technologies) and allowed to cure O/N. Images were acquired on Airyscan microscope (Zeiss) using a 63x objective (oil immersion), 2x pixel sampling and deconvolved using Zen software. Image visualization and analysis were conducted in Fiji^80^.

### Electron microscopy

C1ells (MelJuSo wild type and DKO) were cultured in 10 cm dishes and fixed in half-strength Karnovsky’s fixative (2.5% glutaraldehyde (EMS) and 2% formaldehyde (Sigma)), prepared in 0.1 M PHEM buffer (pH 7.4) at room temperature for 2 hours. After fixation, the samples were rinsed thoroughly with PHEM buffer and stored in 1% formaldehyde at 4°C until further processing. Post-fixation processing was performed in 1% osmium tetroxide (OsO_4_) and 1.5% potassium ferricyanide (K_3_Fe(CN)_6_) in 1 M phosphate buffer (pH 7.4) for 2 hours at room temperature. Samples were then dehydrated through a graded acetone series (70% overnight, followed by 90% for 15 min, 96% for 15 min, and 100% acetone for 3 × 30 min) and embedded in Epon resin (SERVA). Ultrathin sections (70 nm) were cut using a Leica Ultracut UCT ultramicrotome, mounted on formvar- and carbon-coated TEM grids, and stained with uranyl acetate and lead citrate using a Leica AC20 staining system.

Electron micrographs were acquired using either a JEOL JEM1011 transmission electron microscope equipped with a Veleta 2k × 2k CCD camera (EMSIS, Münster, Germany), or a Tecnai12 (FEI Thermo Fisher) microscope, also equipped with a Veleta 2k × 2k CCD camera, operated via SerialEM software. Image visualization and analysis were conducted in Fiji^80^.

### Total internal reflection (TIRF) microscopy

To MVB fusion with the PM, we employed live-cell exosome reporters in which pH-sensitive fluorescent proteins—either super ecliptic pHluorin (green) or pHuji (red)—inserted into the first extracellular loop (EC1) of the EV enriched tetraspanin CD63 ^58^. The acidic pH of late endosomes quenches the fluorescence of these reporters, which is rapidly restored upon fusion with the PM. This results in a burst of fluorescence that can be detected by TIRF microscopy. Fusion events were defined as sudden, transient increases in fluorescence intensity and were quantified as described previously ^56,57^. Fusion activity was expressed as the number of events observed during a standardized time-lapse imaging period—typically 3 minutes—which was kept consistent for each experiment and its replicates. For each condition, an average of ≥10 cells were imaged across ≥5 fields of view in ≥3 independent biological replicates. Image visualization and analysis were conducted in Fiji^80^.

### Pathway enrichment analysis

Gene ontology analysis was performed using DAVID^81^. Analysis of the CRISPRa genetic screen was performed on candidate genes with a combined score > 3. Analysis of biotinylation proteomics was performed on 336 proteins exhibiting greater exclusive unique peptide counts in the MARCKS and MARCKSL1 groups compared to control. All results were visualized using ImageGP^82^.

### SDS–PAGE and western blotting

Protein samples were prepared with Laemmli Sample Buffer. After denaturation, samples were separated by Bolt 10% or 12% SDS-PAGE and transferred onto PVDF or nitrocellulose membrane in ethanol-containing transfer buffer at 300 mA for 1.5-2h. Blots were blocked in 5% milk/PBST for 1Lh, at RT, and probed with the indicated primary antibodies diluted in in 5% milk/PBST overnight, at 4°C. After washing 3 × 10 min in PBS/0.1% Tween-20, proteins were detected with fluorescent (Li-COR Biosciences) or HRP-conjugated secondary antibodies (Invitrogen). Signals were imaged directly using Odyssey Fx laser scanning fluorescence imager (LI-COR Biosciences) or using Chemidoc XRS+ imager (Bio-Rad) immediately following the addition of chemiluminescent substrate (Thermo Fisher Scientific 32209) to the membrane. Quantification of band intensities was performed using LI-COR ImageStudio software or Fiji/ImageJ.

## Statistics

Statistical analyses were performed using GraphPad Prism 10.2.3 as indicated in the corresponding figure legends.

## Data availability

The mass spectrometry proteomics data have been deposited to the ProteomeXchange Consortium via the PRIDE^83^ partner repository with the dataset identifier PXD066258.

## Author contributions

IB, FV and JN designed the study. YZ performed and analyzed most of the experiments. RW designed and performed the CRISPRa screen, with assistance from BC. AG performed TIRF experiments under the supervision of FV. NL and CdH performed EM analysis. RT and GJ carried out mass spectrometry analysis under the supervision of PvV. JA created eML1-GFP cells and aided in genome editing design. MM provided technical support. YZ and IB wrote the manuscript with input from all authors. IB supervised the project. JN, IB and FV contributed financial support.

## Acknowledgements

This work was supported in part by China Scholarship Council (CSC) grant (202103250009) awarded to YZ, a Dutch Ministry of Education, Culture, and Science (OCW) Spinoza grant (00897590) awarded to JN, Dutch Cancer Society grants awarded to IB (14760) and FV (12849), and OCW VIDI grant (213.106) awarded to FV. IB is a principal investigator of the Gravitation Consortium “FLOW” (024.006.036), funded by OCW.

## Disclosure and Competing Interests Statement

The authors declare no conflict of interest.

## Notes

### Competing Interest Statement

The authors have declared no competing interest.

