## Supplementary Legends for "CRISPR activation screen uncovers MARCKSL1 as a gauge for extracellular vesicle secretion"

**Supplementary Figure S1. CRISPR-mediated activation identifies MARCKSL1 as a positive regulator of CD63 surface expression (related to Figure 1).**

**A.** Validation of the top hits from the genome-wide CRISPR activation (CRISPRa) screen for CD63 surface expression. MelJuSo cells stably expressing MS2-p65-HSF1 were transduced with a non-targeting gRNA (gNTC) or individual gRNAs targeting the indicated genes, including MARCKSL1 (gML1_1 and gML1_2). Selected cells were analyzed by flow cytometry against surface exposed CD63. Graph reports on mean fluorescence intensities of CD63, expressed relative to gNTC, from n=3 independent experiments. **B.** Validation of CRISPRa-mediated upregulation of MARCKSL1. Expression of MARCKSL1 transcript (mRNA), analyzed by quantitative PCR (qPCR), in gML1_1 edited MelJuSo cells relative to gNTC. Graph reports on n=4 independent experiments.

**C-E.** Effect of ectopic overexpression of MARCKSL1 on CD63 surface expression. **C.** Immunoblot against MARCKSL1 (ML1) (β-actin as loading control) of MelJuSo ectopically expressing GFP or ML1-GFP. **D.** Flow cytometry analysis of CD63 surface levels in MelJuSo (left) and HeLa (right) cells ectopically expressing GFP versus ML1-GFP. Representative histograms are shown. **E.** Quantification of mean fluorescence intensities expressed relative to paired controls. Graph reports on n=8 (MelJuSo) and n=4 (HeLa) independent experiments.

All bar graphs reflect mean (+/-SD) with statistical significance determined using paired Student *t*-test. *p < 0.05, **p < 0.01, NS: not significant.

**Supplementary Figure S2. Validation of MARCKS/MARCKSL1 ablation (KO) using genomic sequencing (related to Figure 1).**

Genomic sequencing of MARCKS/MARCKSL1 ablation in MelJuSo and HeLa clonal lines. Regions containing the relevant indels introduced with CRISP/Cas9 genome editing are shown.

**Supplementary Figure S3. Analysis of cell surface marker expression as a function of MARCKS/MARCKSL1 loss and rescue (related to Figure 1).**

**A.** Effect of MARCKS/MARCKSL1 ablation on cell surface expression of late compartment and biosynthetic pathway markers. Flow cytometry analysis of wild type (WT) and double knock-out (DKO) MelJuSo cells for cell surface abundance of LAMP1, MHC II, MHC I, and CD71. Representative histograms are shown.

**B-E.** Rescue of MARCKS/MARCKSL1 ablation. Flow cytometry analysis of **B, C.** LAMP1 surface levels in MelJuSo DKO cells or **D, E.** CD63 surface levels in HeLa DKO cells stably transduced with empty vector (GFP), MS-GFP or ML1-GFP. **B, D.** Representative histograms are shown. **C, E.** Quantification of mean fluorescence intensities expressed relative to control DKO cells (GFP) within each group. Bar graphs report on n=5 (MelJuSo) and n=3 (HeLa) independent experiments.

**F-H.** Consequences of MARCKS/MARCKSL1 ablation for EV release. **F.** Particle size profiling of EV isolates from wild type (WT) versus double knock-out (DKO) MelJuSo cells using Nanoparticle Tracking Analysis (NTA). Quantification reflects nanoparticle concentration, relative to control cells (WT) from n=3 independent experiments. **G.** Immunoblot analysis of EV isolates and their corresponding whole cell lysate (WCL) controls derived from WT and ML1 KO cells for EV markers CD63, CD81 and MHC II (β-actin as loading control). **­­­­H.** Quantification of protein abundance in EV isolates expressed as ratio EV / WCL. Graph reports on n=3 independent experiments.

All bar graphs reflect mean (+/-SD) with statistical significance determined using paired Student *t*-test; *p < 0.05, **p < 0.01, NS: not significant.

**Figure S4. MARCKSL1 regulates trafficking of CD63 between endosomes and the PM (related to Figure 4).**

**A.** Intracellular distribution of endogenous MARCKSL1. Representative confocal images of live MelJuSo cells expressing endogenous MARCKSL1-GFP (eML1, green), treated with SiR-lysosome (magenta) to label late endosomes and lysosomes. Scale bars = 10 um.

**B.** Representative confocal image of live MelJuSo cells ectopically expressing wild type ML1-GFP versus mutants ∆ED-GFP or ∆MH2-GFP (green), treated with Lysotracker (magenta). Scale bars = 10 um.

**C, D.** Effect of ED on MARCKSL1-mediated regulation of CD63 cell surface expression. **C.** Representative histograms of CD63 cell surface expression in MelJuSo cells ectopically expressing wild type ML1-GFP versus mutants ∆ED-GFP or ∆MH2-GFP constructs. **D.** Quantification of mean fluorescence intensities expressed relative to control cells (GFP) from n=4 independent experiments.

All bar graphs reflect mean (+/-SD), with statistical significance determined using paired Student *t*-test. *p < 0.05, **p < 0.01, NS: not significant.

**Supplementary Figure S5. Proximity-based proteomics for MARCKSL1 (related to Figure 5).**

**A.** Silver stained SDS-PAGE gel of neutravidin precipitates derived from HEK293T cells transfected with 2HA-TurboID empty vector (TID), MARCKS-TID or MARCKSL1-TID and treated in the presence of exogenous biotin. Visualized samples underwent mass spectrometry analysis.

**B-E.** Validation of candidate interactors of MARCKS (MS) and MARCKSL1 (ML1). Immunoblot analysis of neutravidin precipitates and their corresponding total lysate (TL) inputs from HEK293T cells co-transfected with MS-HA-TID or ML1-HA-TID and Flag-tagged **B.** RDX, **C.** STXBP3, **D.** EHD4 or **E.** STXBP2 and treated with exogenous biotin for the indicated time (min) prior to lysis.

**F, G.** Effects of STXBP3 and Radixin overexpression on CD63 membrane homeostasis. **F.** Representative flow cytometry histograms of CD63 surface expression in HeLa cells, ectopically expressing GFP, GFP-STXBP3, or GFP-RDX. **G.** Quantification of CD63 surface levels, expressed relative to GFP. Graph reports on n=3 independent experiments.

All bar graphs reflect mean (+/-SD) with statistical significance determined using a paired Student *t*-test. **p < 0.01, ***p < 0.001, NS: not significant.

**Supplementary Table S1. CRISPR activation screen identifies MARCKSL1 as a positive regulator of CD63 surface expression (related to Figure 1).** Hit list ranking based on sequencing of gRNA enrichments post-sorting for high CD63 expressing cells. Activating guide RNAs targeting the MARCKSL1 locus are highlighted in yellow.

**Supplementary Table S2. Profiling of EVs in control versus MARCKSL1-overexressing cells by mass spectrometry (related to Figure 3).** Listed are proteins identified in EV isolates from gNTC (batch 1 and batch 2) versus gML1 (batch 1 and 2) MelJuSo cells.

**Supplementary Table S3. Proximity-based TurboID proteomics for MARCKS and MARCKSL1 identify PM anchor proteins and membrane fusion machinery (related Figure 5).** Listed are proteins identified in neutravidin precipitates from HEK293T cells transfected with 2HA-TurboID empty vector (TID, blue), MARCKS-TID (pink) or MARCKSL1-TID (green) and treated in the presence of exogenous biotin. Rankings are compiled relative to empty vector controls.

**Supplementary Movie S1. Intracellular distribution of endogenous MARCKSL1 (related to Figure 4A).** Time-lapse of MelJuSo cells expressing endogenous MARCKSL1-GFP (green) and treated with SiR-Lysosome (magenta) to label endolysosomes (9 s at 5 frame/sec). Scale bar: 10 μm.

**Supplementary Movies S2. MVB—PM fusion as a function of MARCKSL1 expression (related Figure 4G).** TIRF imaging of exogenously expressed CD63-pHluorin in **A-D.** wild type (WT) or **E.** MARCKS/MARCKSL1 double-knockout DKO cells MelJuSo cells cotransfected with **A.** GFP empty vector, **B.** ML1-GFP, or **C.** ∆ED-GFP. Bright spots appearing at the PM reflect individual MVB—PM fusion events. Displayed at 4x normal speed.
