## Supplementary figures and images for "CRISPR activation screen uncovers MARCKSL1 as a gauge for extracellular vesicle secretion"

### Supplementary Figure S1

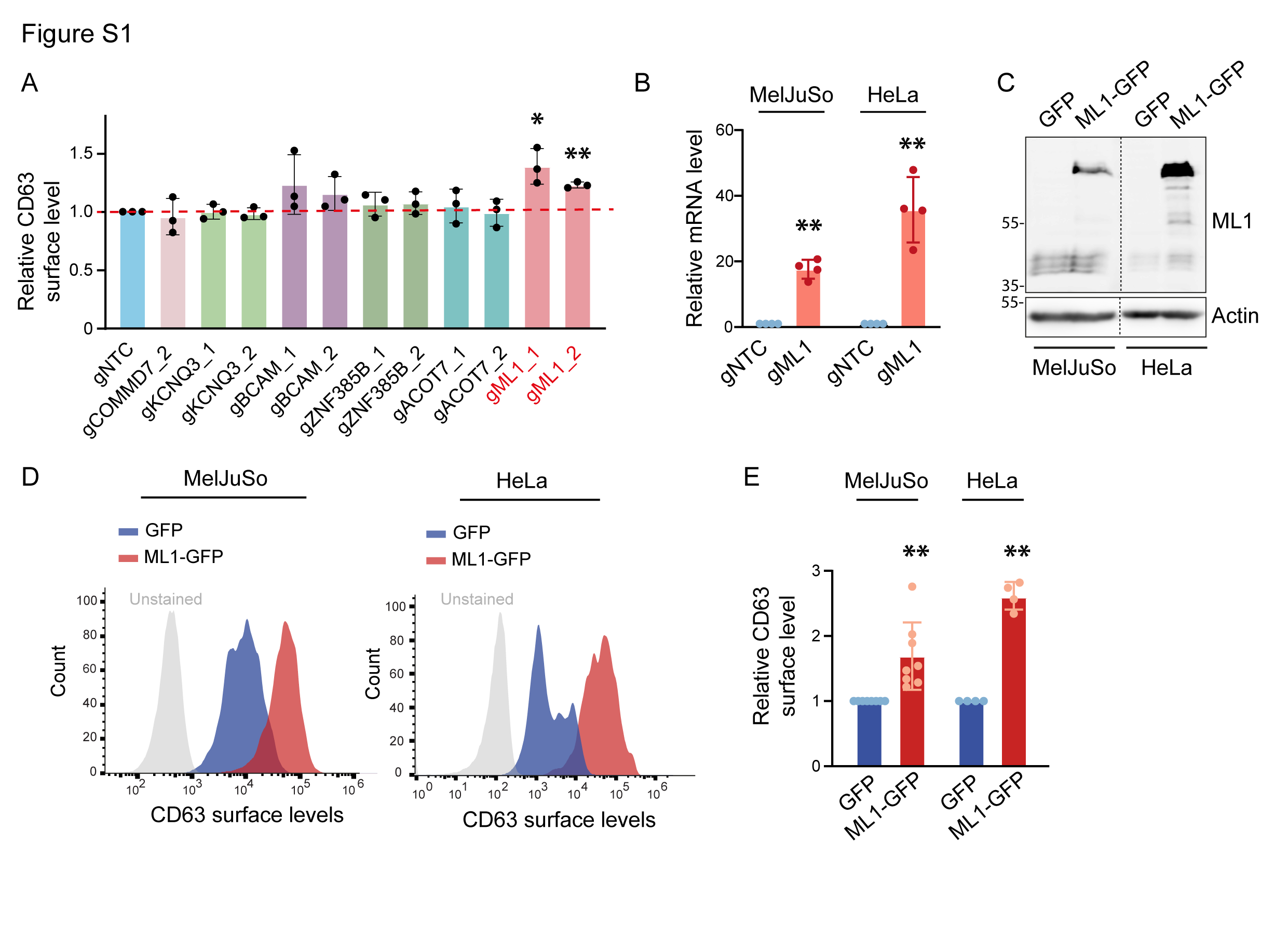

### Supplementary Figure S2

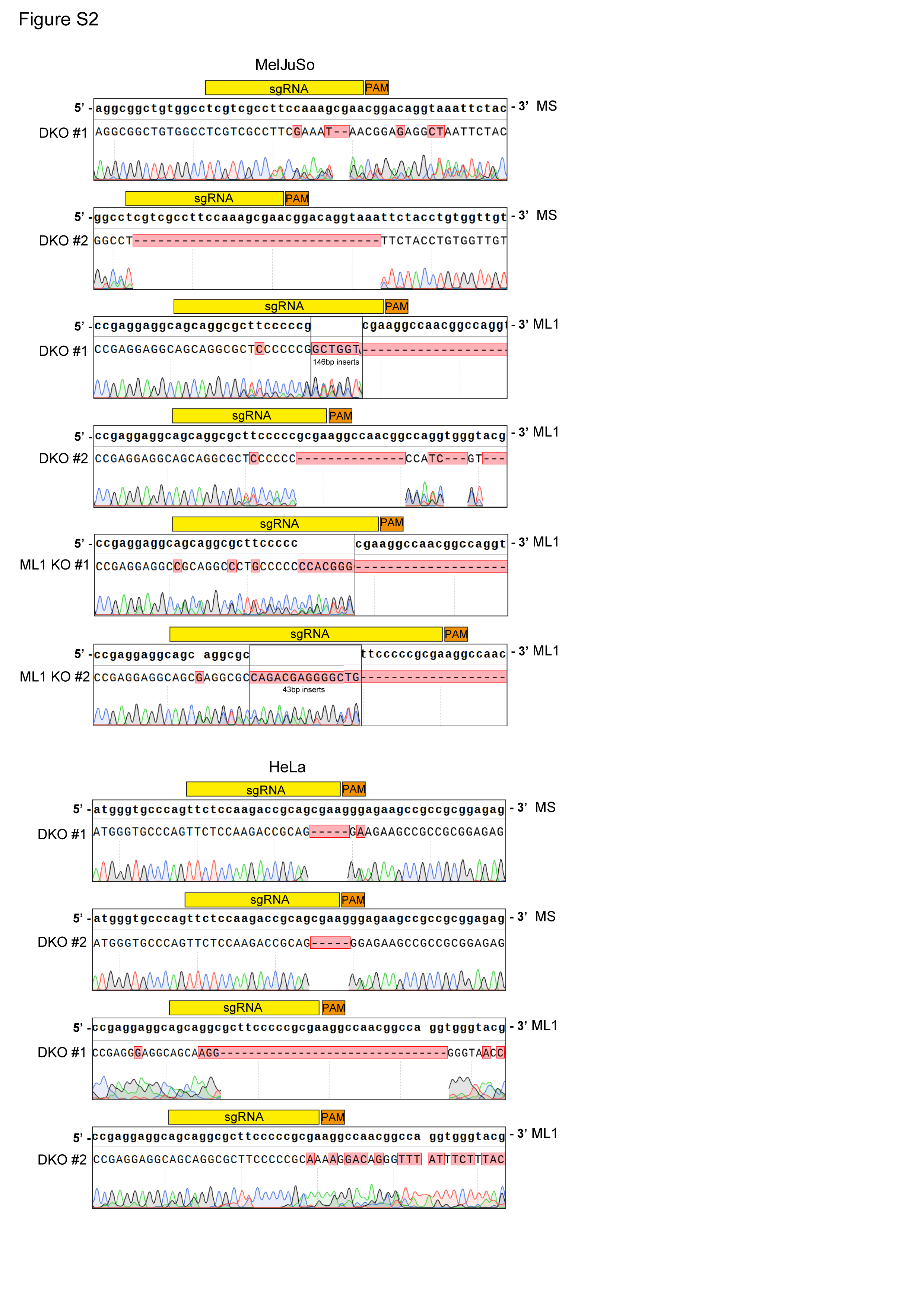

### Supplementary Figure S3

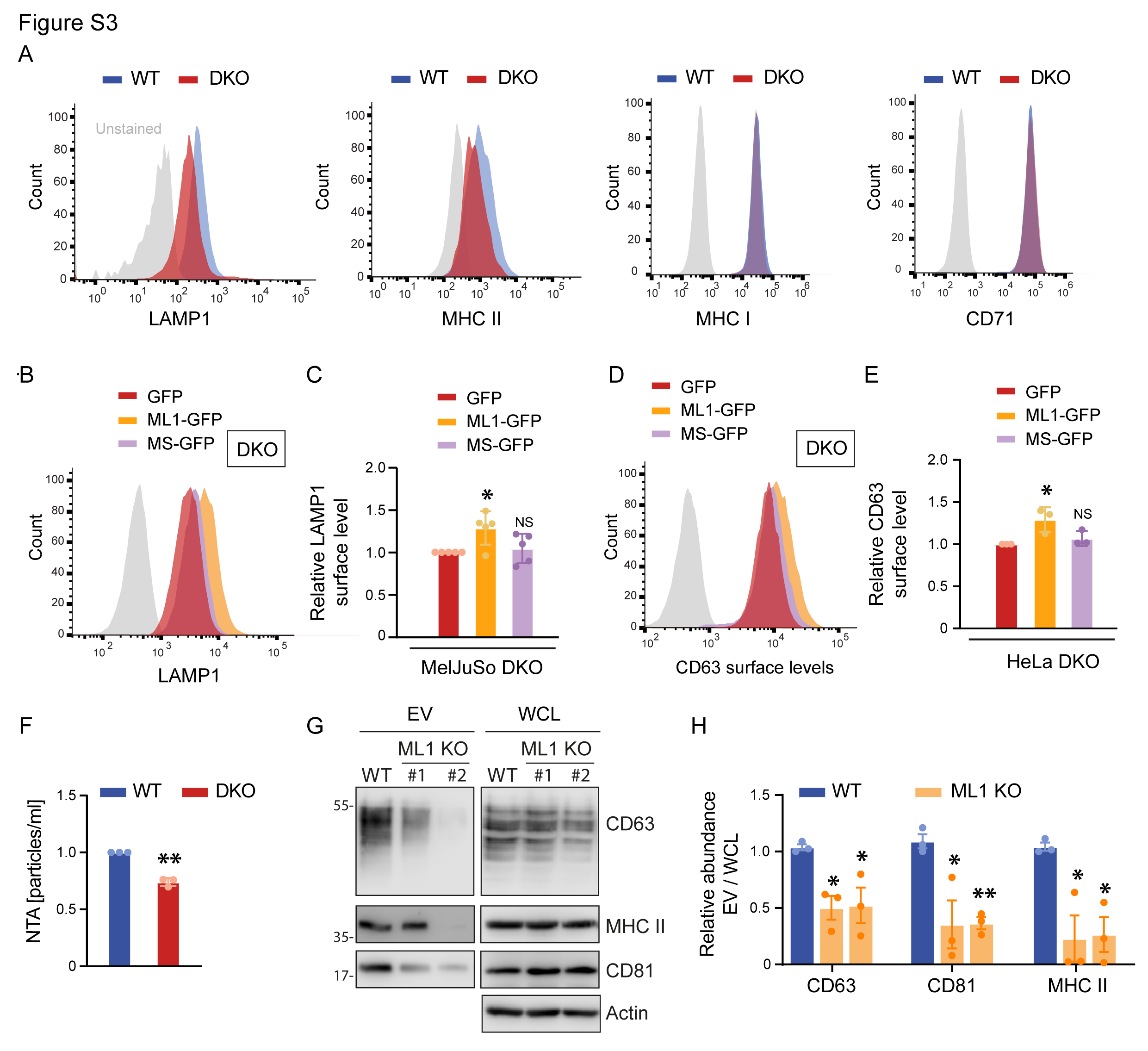

### Supplementary Figure S4

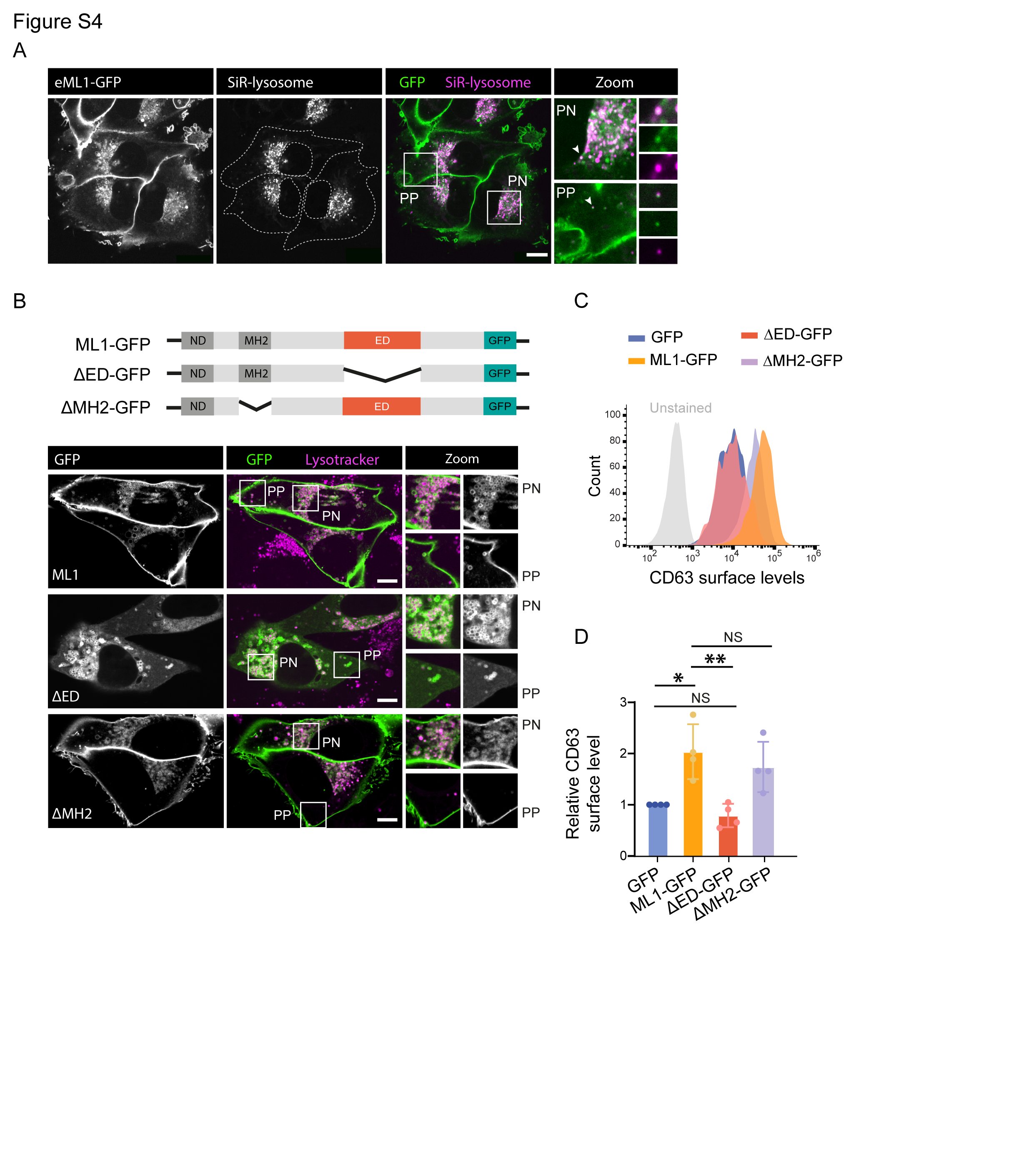

### Supplementary Figure S5

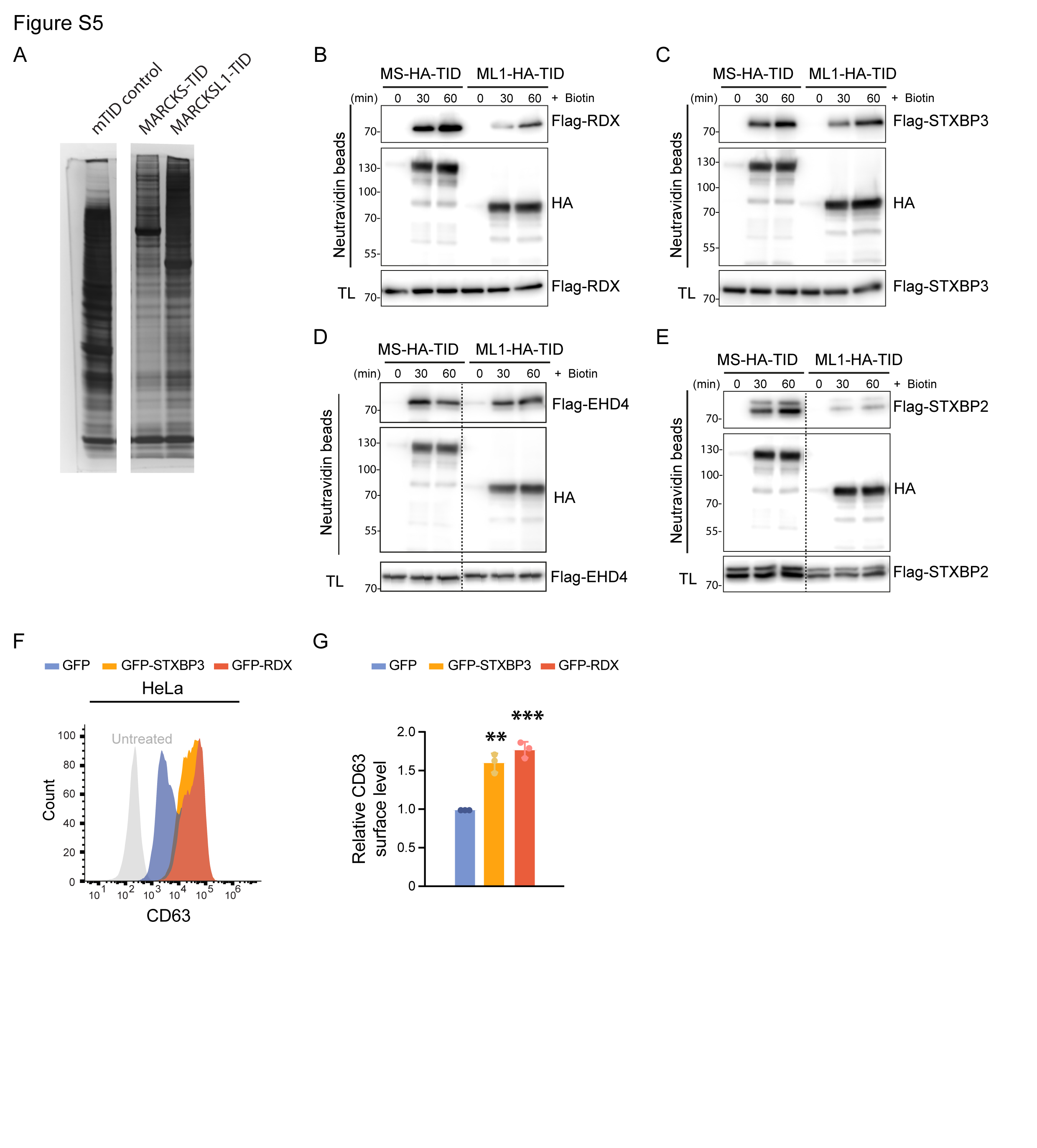
